# Molecular Mechanisms of Chinese Medicine in Regulating Colorectal Cancer Immune Microenvironment: Insights from Single-Cell Transcriptomics and Network Pharmacology

**DOI:** 10.1101/2025.03.24.644909

**Authors:** Chenlong Zhang, Yumei Zhang, Pengfei Li, Yujie Wang, Kaihang Guo, Chunfang Zhang

## Abstract

**Background:** The molecular mechanisms underlying the efficacy of Traditional Chinese Medicine (TCM) in colorectal cancer treatment remain largely unexplored. We developed a computational systems biology approach integrating single-cell transcriptomics with network pharmacology to elucidate the potential mechanisms of TCM in modulating colorectal cancer progression.

**Methods:** We developed an integrated computational pipeline for multi-omics data analysis combining single-cell transcriptomics with network pharmacology. Raw single-cell RNA-seq data from 3 normal tissues and 3 colorectal tumors were obtained from GEO database and processed using a customized workflow in R. Quality control, normalization, and dimensionality reduction were performed using the Seurat v4.0 algorithm, followed by unsupervised clustering to identify cell subpopulations. Differentially expressed genes (DEGs) were identified using MAST algorithm with adjusted p-value < 0.05 and |log_2_FC| > 1.0. These computationally identified DEGs were subsequently mapped to a comprehensive Traditional Chinese Medicine (TCM) database using a network pharmacology approach to predict herb-target interactions. In parallel, we integrated TCGA RNA-seq data (STAR-counts) with clinical information, applying log_2_(TPM+1) transformation for normalization. We then implemented a machine learning-based correlation analysis to construct gene-cell-immunity-pathway networks, using weighted gene co-expression network analysis (WGCNA) to identify key regulatory modules.

**Results:** Our computational analysis of single-cell RNA-seq data identified 109 differentially expressed genes (DEGs) that define the molecular signature of colorectal cancer microenvironment. Clustering algorithms revealed 14 distinct cell subpopulations, with predominant immune cell infiltration, particularly B and T lymphocytes, suggesting a complex immune regulatory network. Network pharmacology analysis mapped these DEGs to potential therapeutic targets, computationally predicting interactions with 140 traditional Chinese herbs. These herbs were classified into 8 functional categories. Through integrative multi-omics analysis and pathway enrichment algorithms, we identified core regulatory networks comprising 23 genes and 39 significantly enriched signaling pathways (FDR < 0.01) that orchestrate immune cell function in the tumor microenvironment. Notably, our analysis in silico revealed previously uncharacterized gene-pathway interactions that may explain the immunomodulatory effects of specific herbal compounds.

**Conclusions:** Our systems biology and computational analysis revealed a potential mechanism by which 8 categories of Chinese herbal medicines and 23 genes across 39 signaling pathways may regulate colorectal cancer progression through modulation of specific gene regulatory networks and immune cell functions. These findings demonstrate the value of integrative computational approaches in elucidating complex biological mechanisms of traditional medicines

---

Colorectal cancer (CRC) is one of the most common malignancies in clinical life. Due to objective and subjective factors, most patients are diagnosed with advanced stages of the disease and the number of cases is gradually increasing, resulting in the third and second highest incidence and mortality rates in the world respectively ^(1)^. Although new advances such as targeted therapies and immunotherapy have advanced by leaps and bounds, chemoradiotherapy remains the mainstay of treatment for advanced colorectal cancer and is not as effective as it could be ^(2)^. Traditional Chinese medicine(TCM), a non-negligible treatment method, plays an indelible role in preventing and treating Colorectal cancer^(3)^. Its advantages in treating Colorectal cancer are to improve the tumor growth environment, stabilize the tumor, improve the patient’s quality of life, reduce the adverse effects of chemoradiotherapy, and prolong the survival period ^(4)^. A retrospective cohort study on colorectal cancer was investigated the survival between 147 patients of Chinese herbal medicine (CHM) users and 388 patients of CHM nonusers, the multivariate Cox regression analysis showed that CHM use was significantly associated with better survival^(5)^. Meanwhile, a randomized, double-blind, placebo-controlled clinical trial was collected 200 first-line and 120 second-line patients with Metastatic Colorectal Cancer who underwent chemotherapy combined with Cetuximab or Bevacizumab. The treatment group was combined with Huangci Granule(a traditional Chinese medicine for treating metastatic colorectal cancer), while the control group was combined with placebo. The result showed that the median progression-free survival (PFS) was 9.59 months *vs* 6.89 months in the treatment group and control group in the first-line treatment. Chinese medicine was an independent factor affecting the PFS. In the second-line treatment, the median PFS was 6.51 months *vs* 4.53 months in the treatment group and control group. Compared with the control group, role function, social function, fatigue and appetite loss were significantly improved in the treatment and drug related grades 3 to 4 adverse events were less^(6)^.

Despite the indelible contribution of TCM in treating patients with colorectal cancer, the complex composition of TCM generates complex physiological processes in the body, which makes it difficult for us to unravel its mechanism^(7)^; moreover, the occurrence and development of colorectal cancer are regulated by complex network pathways in the body, which further makes it difficult for us to elucidate the targets of action of TCM in treating patients with tumors.

In recent years, novel single-cell sequencing technologies have allowed for the clustering of cells and facilitated their classification. The clustering of transcriptomes facilitates the automatic classification of cells and enables the identification of heterogeneous cell types and molecular states, while the sequencing of single-cell genomes or transcriptomes to reveal cell population differences and cellular evolutionary relationships can reveal the complex heterogeneous mechanisms involved in disease onset and progression^(8)^. At the same time, the gradual improvement and multi-functionality of the database of Chinese medicine can reveal the active ingredients and targets of action of Chinese medicine^(9, 10)^. It is essential to establish a correlation between single-cell sequencing technology and TCM databases to analyze the mechanism of action of TCM in treating patients with colorectal cancer.

## Methods

### Data Screening

We obtained the raw data of single-cell transcriptome profiling(Tissue: colorectal cancer, Data Set Number: GSE163974, Chip Platform: GPL16791, Sample Type: 3 normals and 3 tumors, Species: human, Data Types: 10x Genomics) from the Gene Expression Omnibus (GEO) database (https://www.ncbi.nlm.nih.gov/geo/). We utilized the Seurat package in R studio to generate the object and filter out cells with poor quality. Then, we conducted standard data preprocessing, where we calculated the percentage of the gene numbers, cell counts, and mitochondria sequencing count. We excluded genes with less than 10 cells detected and disregarded cells with less than 200 detected gene numbers.

### Data Preprocessing

Only genes detected in at least 10 cells were retained. We filtered out cells with more than 500 detected genes and count RNA(1,000<n<20,000), and those with a high mitochondrial content (>95%). After discarding poor-quality cells, a total of 485 cells were retained for downstream analysis. To normalize the library size effect in each cell, we scaled UMI(Unique Molecular Identifiers)counts using scale. factor = 10,000. Following the log transformation of the data, other factors, including “percent. mt”, “nCount_RNA” and “nFeature_RNA”, were corrected for variation regression using the ScaleData function in Seurat (v4.3.0). See the flow chart(Figure 1) for details of the process.

**Figure 1.**
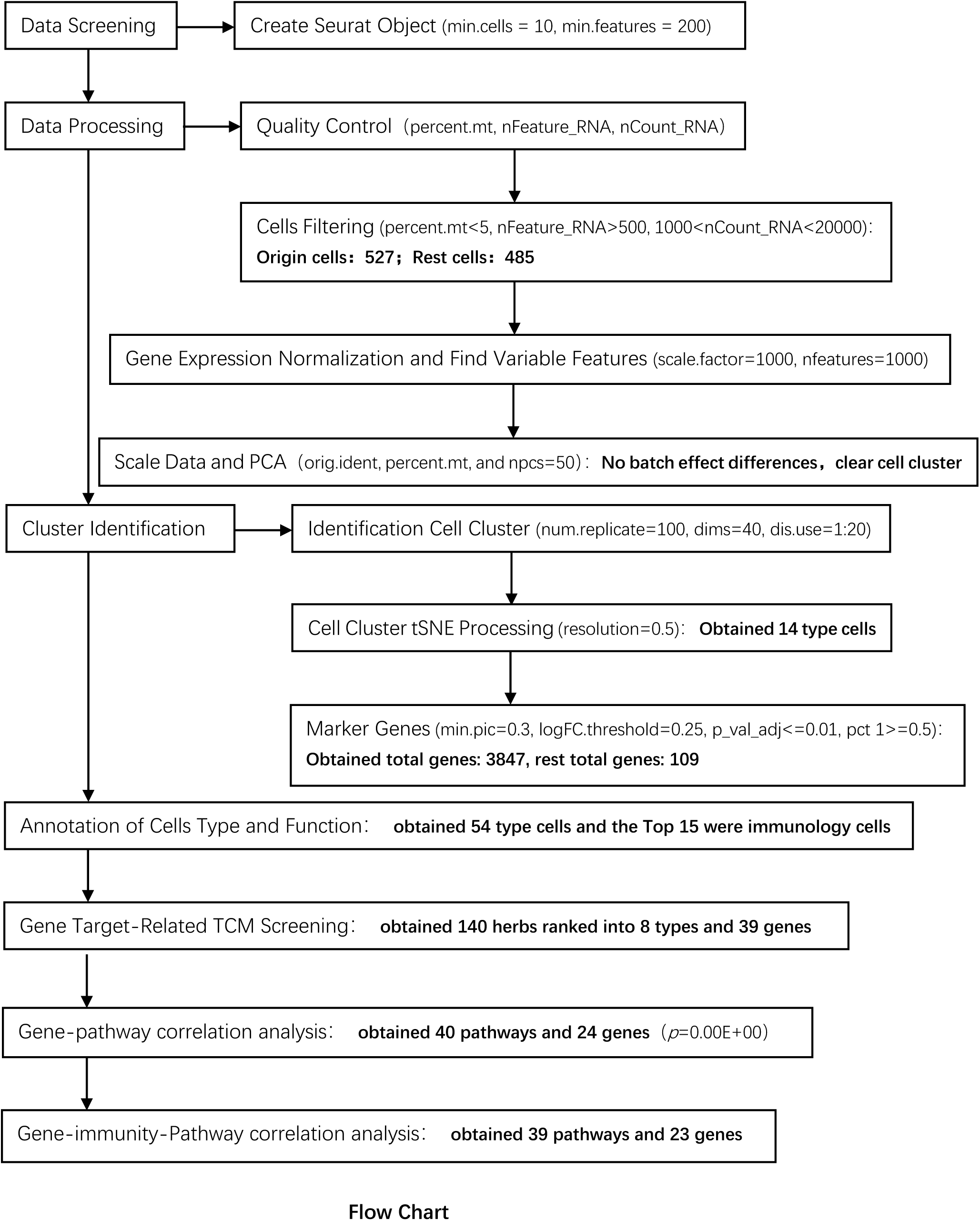
Flow Chart. This flow chart succinctly represents the entire analysis process and the results of each step, where the downward-facing boxes and arrows on the left demonstrate the main analysis steps, and the boxes and downward-facing arrows on the right represent the conditions and results obtained at each step.

### Cluster Identification

The corrected normalized data metrics were applied to the standard analysis as described in the Seurat R package. The top 109 variable genes were extracted for principal component analysis (PCA). The top 20 principal components were kept for tSNE visualization and clustering. We performed cell clustering using the FindClusters function (resolution =0.5) implemented in the Seurat R package (MAST algorithm with adjusted p-value < 0.05 and |log2FC| > 1.0). See the flow chart(Figure 1) for details of the process.

### Annotation of Cell Type and Function

The Human CellMark dataset was downloaded from the website (http://biocc.hrbmu.edu.cn/CellMarker/download.jsp) and loaded into the R Studio platform. Matches between the Human CellMark and the clusters associated with colorectal cancer were made by gene name, annotated for cell type and function in cluster identification, and manually checked against the literature step by step. Accuracy was checked and non-colorectal cancer tissue-associated cells were manually excluded. See the flow chart(Figure 1) for details of the process.

### Gene Target-Related Chinese Medicine Screening

The obtained genes above as targets were searched separately on the website(herb.ac.cn/) to obtain the names of herbal medicines and the accuracy was determined in the literature.

### Gene-pathway correlation analysis by weighted gene co-expression network analysis (WGCNA)

We downloaded STAR-counts data and corresponding clinical information for colorectal cancer from the TCGA database (https://portal.gdc.cancer.gov). We then extracted data in TPM format and performed normalization using the log2(TPM+1) transformation. After retaining samples that included both RNAseq data and clinical information, we ultimately selected 620 samples for further analysis. We collected the 39 genes included in the corresponding 105 pathways and then analyzed them using the GSVA package in R software, choosing the parameter method= ‘ssgsea’for single-sample gene set enrichment analysis (ssGSEA). Finally, we studied the correlation between gene expression and pathway scores through Spearman correlation analysis. Statistical analysis was conducted using R software, version v4.0.3. Results were considered statistically significant when the p-value was less than 0.05. The pathways corresponding to the 39 genes were sorted in reverse order from smallest to largest based on the p-value, and then filtered based on the p-value equal to 0.00E+00.

### Gene-immunity correlation analysis through six algorithms

We downloaded STAR-counts data and corresponding clinical information for colorectal tumors from the TCGA database (https://portal.gdc.cancer.gov). We then extracted data in TPM format and performed normalization using the log2(TPM+1) transformation. After retaining samples that included both RNAseq data and clinical information, we ultimately selected 620 samples for further analysis. We will use Spearman’s correlation analysis to measure the monotonic relationship between quantitative variables that do not follow a normal distribution. Statistical analysis was conducted using R software, version v4.0.3. Results were considered statistically significant when the p-value was less than 0.05. In addition, it is added here that the data related to CLUL gene were not included in the above databases, so there was no analysis related to CLUL gene in this section.

To conduct a reliable immune score assessment, we used immunedeconv, an R package that integrates six state-of-the-art algorithms, including TIMER, xCell, MCP-counter (Microenvironment Cell Populations-counter), CIBERSORT (Cell-type Identification By Estimating Relative Subsets Of RNA Transcripts), EPIC (Estimating the Proportion of Immune and Cancer cells), and quanTIseq (quantitative deconvolution of RNA-seq). These algorithms have been systematically benchmarked, and each has shown its unique performance and advantages. We used the R package ggClusterNet for analysis and visualization of the results. Tumor Immunophenotype (TIP) is a system that integrates two existing methods, “ssGSEA” and “CIBERSORT”, to track and analyze the proportion of tumor-infiltrating immune cells in the cancer immunity cycle. All the above analysis methods and R packages were performed using R software version v4.1.3. *p* < 0.05 was considered statistically significant.

## Results

### High-Quality Data

To improve data quality and exclude low-quality data, we used percent.mt, nFeature_RNA, and nCount_RNA as metrics for quality control. As shown in the violin diagram, in nFeature_RNA, the gene expression in normal tissues was below 2000, while that in colorectal cancer tissues was below 4000; in nFeature_RNA, the RNA expression in normal tissues was below 500, while that in colorectal cancer tissues was around 2000; in percent.mt, the mitochondria in normal tissues mostly in 5% and in tumor tissues mostly in about 10%. This indicates that this experiment data is of high quality and can be used for further analysis and research. In the FeatureScatter plot(Figure 2), nFeature_RNA and nCount_RNA for normal and tumor tissues are almost linearly positively correlated, while the percentage of mitochondria is negatively correlated with increasing nCount_RNA, which indicates the high quality of this data and the absence of a large number of mitochondrial genes involved in the subsequent analysis, as well as the absence of significant loss and corruption of the data authenticity during the quality control process.

**Figure 2.**
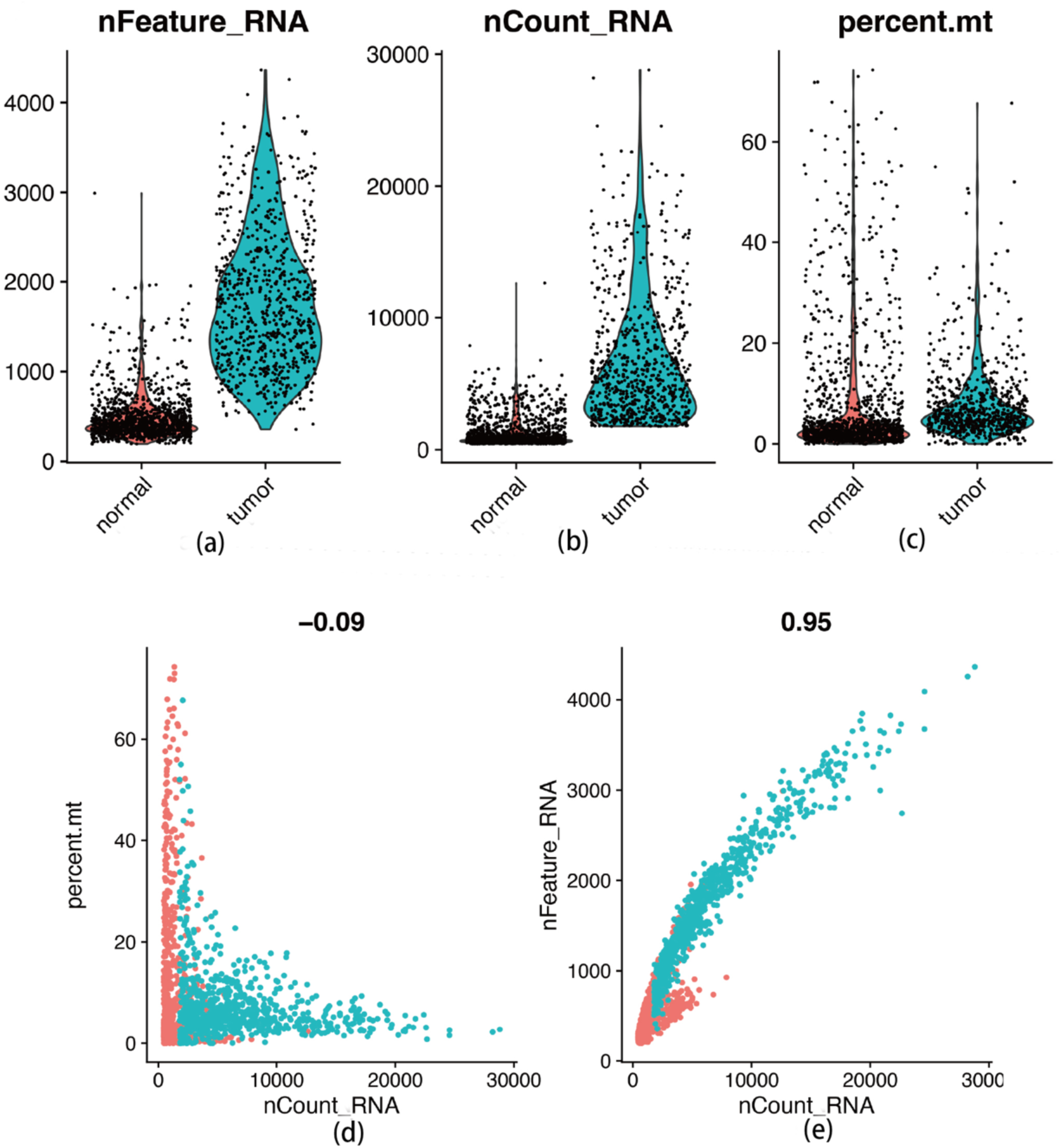
Quality Control Plot. (a) Quality Control Plot-violin plot. The left graph shows the distribution of the number of genes identified in each cell of normal and colorectal cancer tissues. (b) Quality Control Plot-violin plot. The middle graph displays respectively the distribution of the number of RNAs between the two groups. (c) Quality Control Plot-violin plot. The right graph displays the distribution of the mitochondria in each of the two groups as a percentage of the total number of RNAs in each of the two groups. This QC plot can facilitate us to understand the basic data distribution of this single-cell sequencing and help us carry out the subsequent cell screening process. (d) Quality Control Plot-scatter plot. A scatter plot of the number of nCount_RNA in each cell versus the corresponding percentage of mitochondria identified is shown, from which it is clear that there is a strong negative correlation between the two variables, this indicates that as the sequencing volume increases, the number of mitochondria that can be identified in each cell is also decreasing accordingly. (e) Quality Control Plot-scatter plot. A scatter plot of the number of nCount_RNA in each cell versus the corresponding number of genes identified is shown, from which it is clear that there is a strong positive correlation between the two variables. This indicates that as the sequencing volume increases, the number of genes that can be identified in each cell is also increasing accordingly.

### High-Purity Cells through Cells Filtering

Cell purification helps to improve the quality and accuracy of the sequencing data by reducing variability introduced by heterogeneous cell populations. This enables more precise identification of cellular heterogeneity and gene expression patterns within the analyzed sample. For the classification of cells and analysis of differential genes in subsequent studies, the cells were filtered under the conditions (percent. mt<5, nFeature_RNA>500, 1000<nCount_RNA<20,000), and the final results showed: origin cells: 527; remaining cells: 485. The main stringent consideration for filtering cells is to keep cells high quality, with mitochondrial genes accounting for less than 10%; the number of genes expressed should be at least 500; The number of RNA molecules measured should be neither less than 1,000 nor more than 20,000; if the number of RNA molecule fractions is too less, it indicates the cells are insufficient, and if it is much more, it indicates cellular impurity. More details are shown in Figure 3.

**Figure 3.**
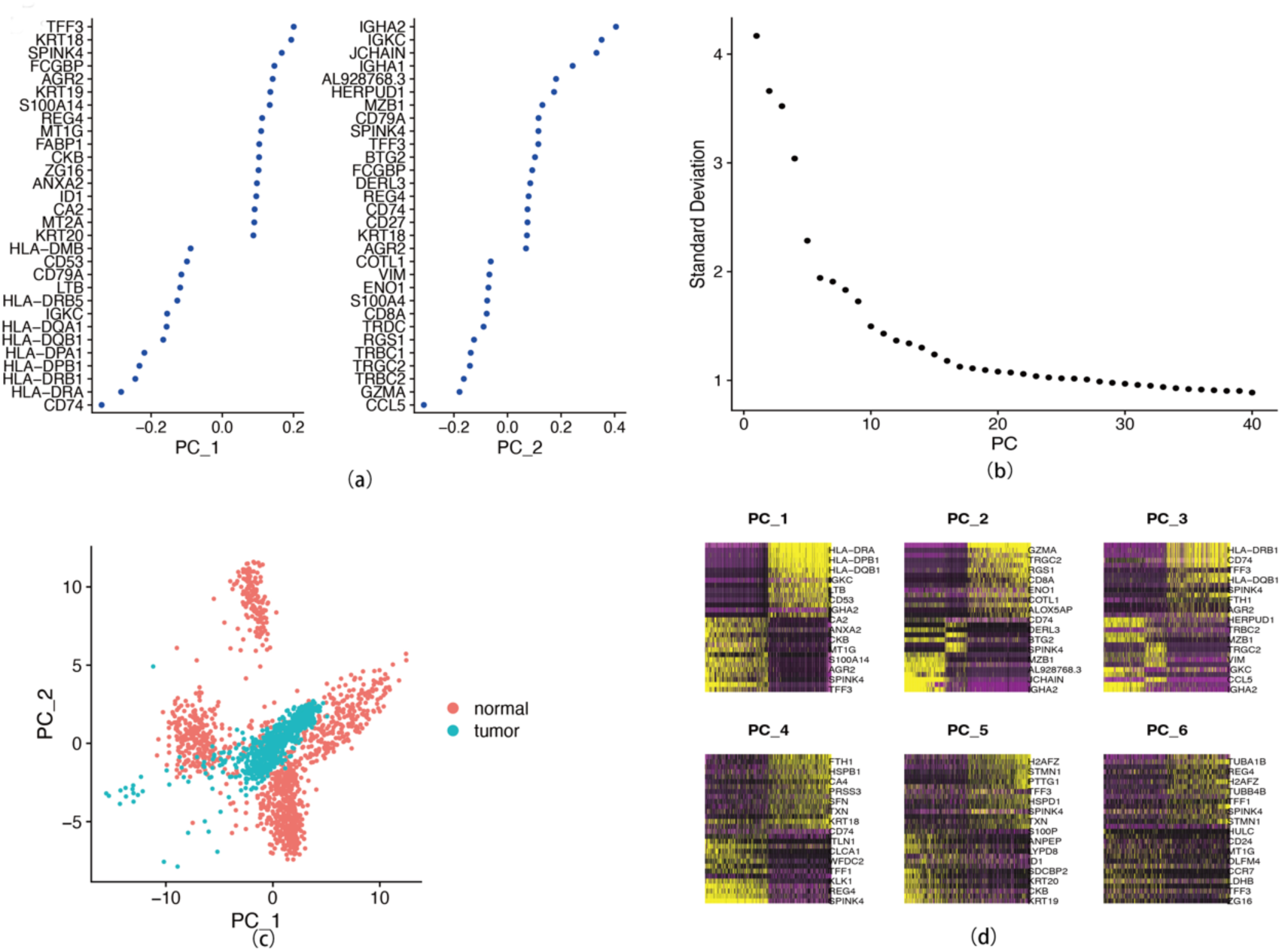
Principle Component Analysis (PCA)Plot. (a) Visualize a Dimensional Plot. The differential genes obtained from our analysis were grouped in cohorts of every 50 genes, with each point representing 50 genes with similar partitioning, respectively. (b) Elbow Plot. As the curve declines to a horizontal coordinate value of 20, the subsequent curve approximates a straight line, suggesting that 20 is the inflection point and that its subsequent genes do not vary much in their differences among individual cells. (c) Dimplot. Red dots represent normal tissue cells and green dots represent colorectal cancer cells. (d) Dimheatmap. The first figure separates the cells into two categories, and the other subsequent figures are increasingly unable to distinguish between the two categories of cells clearly.

### Gene Expression Normalization and Variable Features

To remove technical biases and variations introduced during the scRNA-seq experiment, ensuring that gene expression levels can be accurately compared between cells and appropriately obtained the variable features, the gene normalization was execuated under the condition of scale. factor = 10000 and nfeatures = 2000, the LogNormalize algorithm was used to normalize gene expression and to find differential genes, as shown in Figure 4, with a linear positive correlation between the number of differential genes as gene expression increased. And the genes with significant differences were shown such as Top 10: IGLC2, ZG16, IGHA1, REG1A, IGLL5, JCHAIN, IGLC3, C1QA, IGFBP7, and CA4. For IGLC2, also known as Immunoglobulin lambda constant 2, is a human gene that encodes for one of the constant regions of the immunoglobulin lambda light chain. Immunoglobulin lambda chains are components of antibodies, which are crucial for the immune system’s ability to recognize and neutralize foreign invaders like bacteria and viruses. For IGHA1, one of the genes within the IGH family, it encodes the alpha chain of IgA1. IgA1 is a subclass of immunoglobulin A, one of the main classes of antibodies found in the human body. IgA1 antibodies are primarily found in mucosal secretions, such as saliva, tears, and mucous membranes of the respiratory, gastrointestinal, and genitourinary tracts. The IGLL5 gene, also known as Immunoglobulin lambda-like polypeptide 5, is a gene that encodes for a protein involved in the development and function of B cells, which are a type of white blood cell that produces antibodies as part of the immune response. The JCHAIN gene, also known as joining (J) chain, is a human gene that encodes a protein called the J chain. This gene is primarily expressed in B lymphocytes, where it plays a crucial role in the production and secretion of polymeric immunoglobulins, such as IgA and IgM. It is also involved in the assembly and transport of polymeric immunoglobulins across mucosal surfaces and into secretions, such as saliva, tears, and mucous membranes of the respiratory, gastrointestinal, and genitourinary tracts. It mediates the association of individual antibody monomers, such as IgA or IgM, into polymeric structures, thereby enhancing their stability and functionality in mucosal immunity. In addition to its role in antibody polymerization, the J chain also contributes to the interaction of polymeric immunoglobulins with mucosal epithelial cells and other components of the mucosal immune system, facilitating immune defense against pathogens at mucosal surfaces. IGLC3, immunoglobulin lambda constant 3, a human gene that encodes for one of the constant regions of the immunoglobulin lambda light chain, is similar with IGLC2. The C1QA gene is a human gene that encodes for the A chain of the complement component 1, q subcomponent (C1q). C1q is a protein complex involved in the classical pathway of the complement system, which is a crucial part of the immune system responsible for identifying and eliminating pathogens, as well as clearing damaged cells and immune complexes. In this top10 genes, most of the genes are related to immunity, which plays a crucial role in drawing our conclusions.

**Figure 4.**
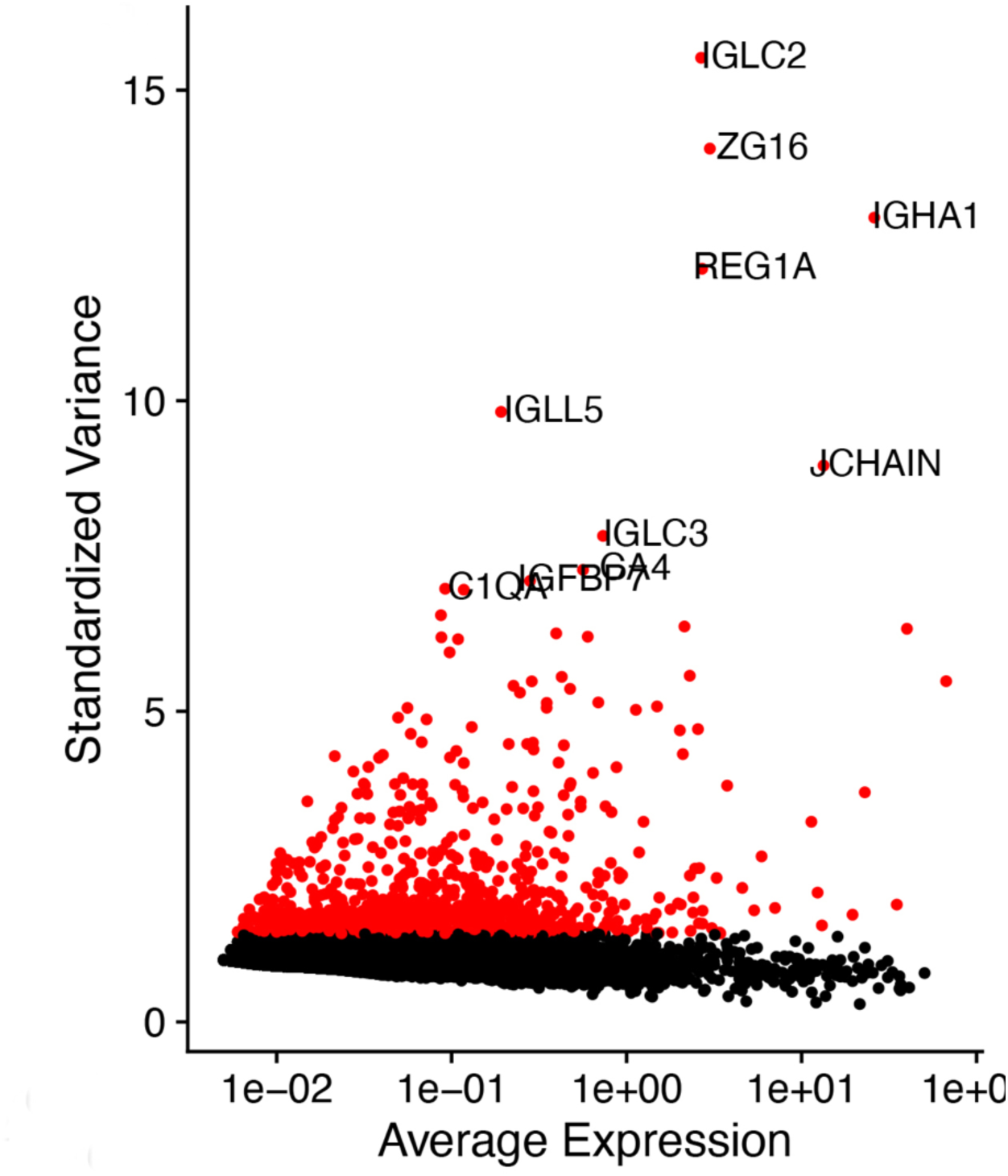
Cellular Differential Gene Distribution Plot. Distribution of differential genes between cells after gene LogNormalization; standardized variance expression gradually increased with increasing average gene expression. This indicates that we obtained high-quality data to ensure the accuracy and correctness of the subsequent analysis.

### No Batch Effect Differences and Clear Cell Clusters

To eliminate batch effects and the effect of mitochondrial number on the overall data, homogenization was manipulated under the condition (orig. ident, percent. mt, and npcs=50, npcs means all analyzed genes were packaged into 50). To determine the presence or absence of batch effects and perform cell clustering, the principal components analysis (PCA) was used. According to Figure 5, no batch effect differences were found, and the cell cluster was clear. This process ensures a high-quality computational procedure for the subsequent accurate and precise acquisition of cell clustering and annotation of cell type names and functions.

**Figure 5.**
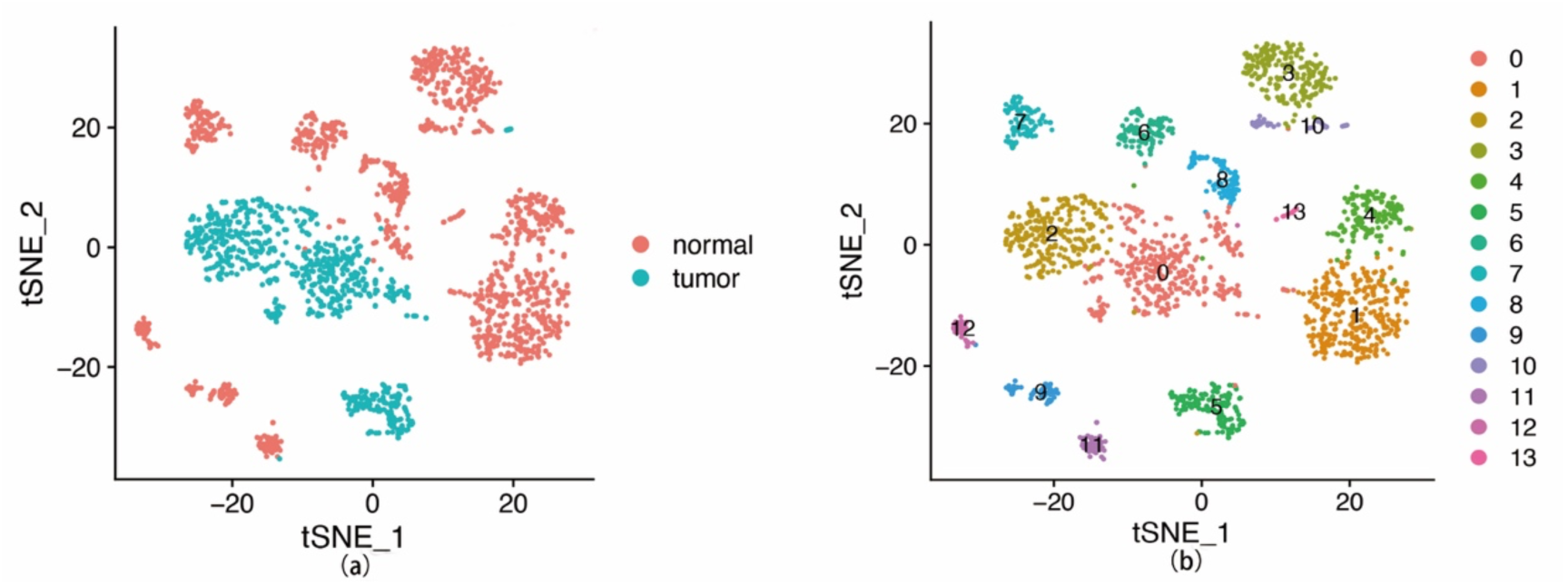
Cell Clustering Analysis Plot. (a) tSNE analysis for normal and tumor. It is shown that cells were divided into tumor and normal cells. (b) tSNE analysis for normal and tumor cell clustering. It is shown that cells were divided into 14 types.

### 14 Categories of Mostly Immune-Related Cells

The tSNE clustering algorithm was used for cell clustering analysis under the condition of dim.use=1:20 which was determined by running the ScoreJackStraw function and resolution=0.5. 14 classes of cells are shown in Figure 5. After annotation of the classified cells, it was found that most of these 14 types of cells were cells with immune functions as shown in Table 1. The top cells were ranked according to the number of target genes in the cells, and the following cells were obtained: B cel(l 18 genes), Cancer stem cell (17 genes), liver stem cell (17 genes), Epithelial cell (12 genes), T cell (12 genes), Cancer cell(11 genes), Macrophage (6 genes), Basal cell (5 genes), Eosinophil (4 genes), Oligodendrocyte(4 genes), Circulating fetal cell(4 genes), Lymphocyte(4 genes), Natural killer cel(l 3 genes), Mesenchymal stromal cell(3 genes), Placenta cel(l 3 genes). Most of them were immune cells after categorization and analysis. Combined with the above obtained differential genes, we can clarify which cells and which differential genes affect the development of colorectal cancer, which will help us to identify the specific target genes and target cells for the action of TCM, and help us to explain the possible mechanism of action of TCM.

**Table 1.**
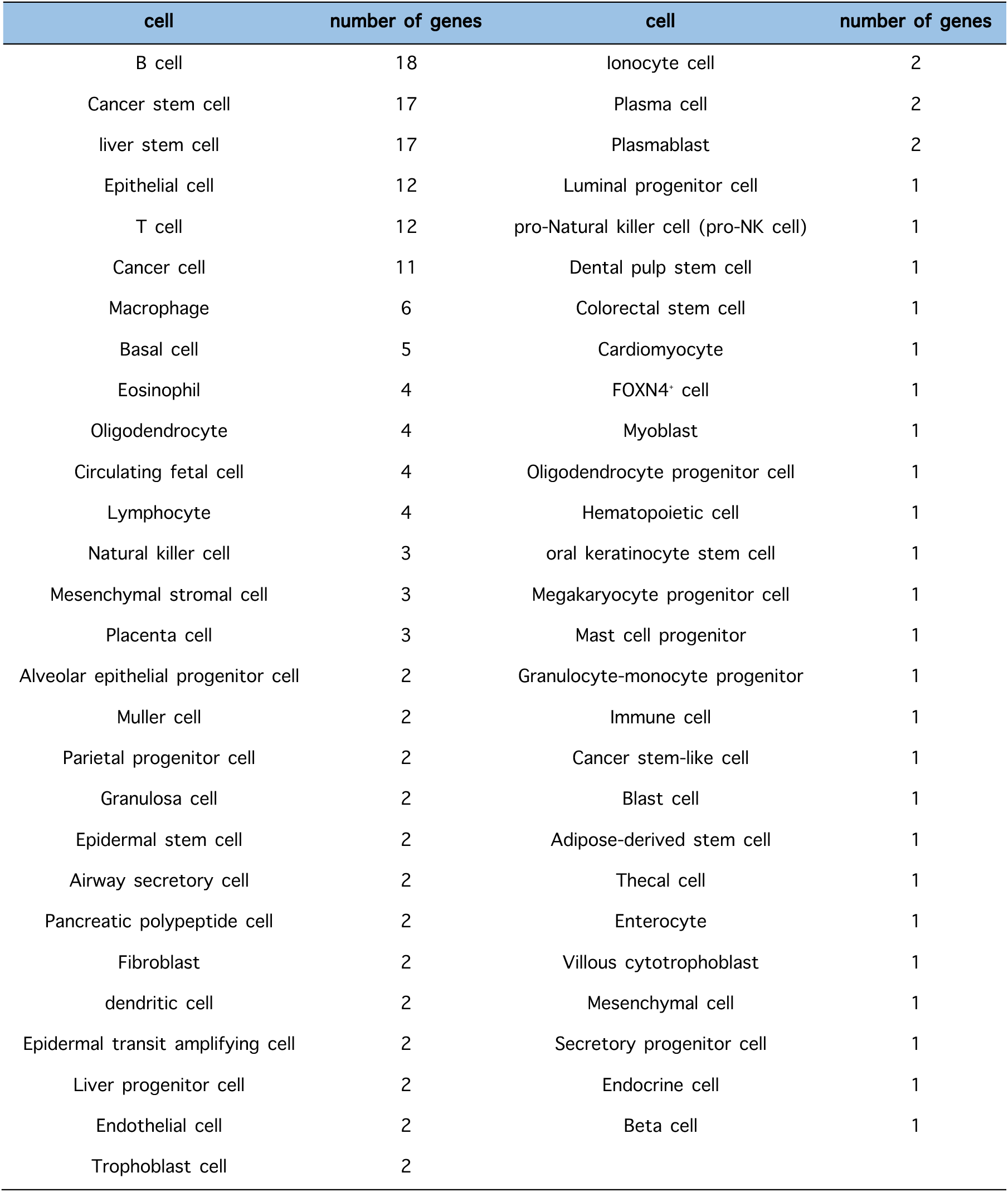
The Classified Cells after Annotation and the Associated Number of Target Genes.

### 8 Types of Chinese Medicines Might Affect on Differential Genes in 14 Types of Immune-Related Cells

The target genes were obtained through the analysis of single-cell sequencing data, and 8 categories of Chinese herbal were retrieved through the Chinese Medicine database, which is mainly focused on certain aspects such as clearing heat, clearing away toxins, relieving dampness, removing blood stasis, strengthening the spleen, benefiting the kidney, tonifying Qi, and nourishing the blood as the Table 2 shown. This database named HERB, a high-throughput experiment-and reference-guided database of Traditional Chinese Medicine, is providing 4 types search items, including: herb, ingredient, target and disease. In this study, we searched all genes in Figure 4 and obtained 140 types herbs and then classified into 8 type of herbal medicines according to the function in TCM. Reviewing the whole research process, we started from single-cell sequencing analysis, obtained differential genes and cell clustering, attempted to functionally annotate the obtained clustered cells, used the differential genes as the targets to reverse search for the names of Chinese herbs in the database, and ultimately obtained the 8 classes of Chinese herbs compatible with clinical treatment of colorectal cancer.

**Table 2.**
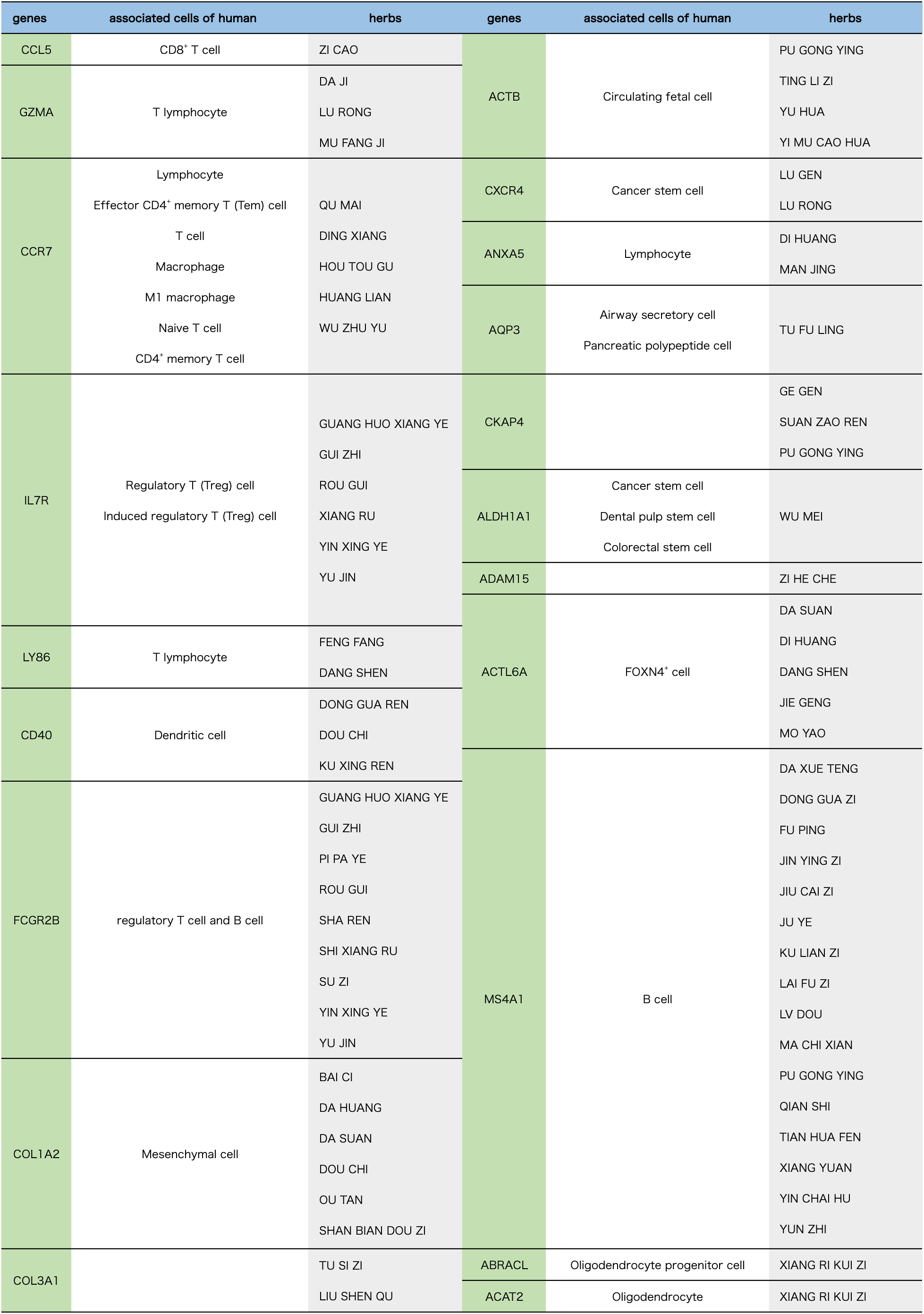
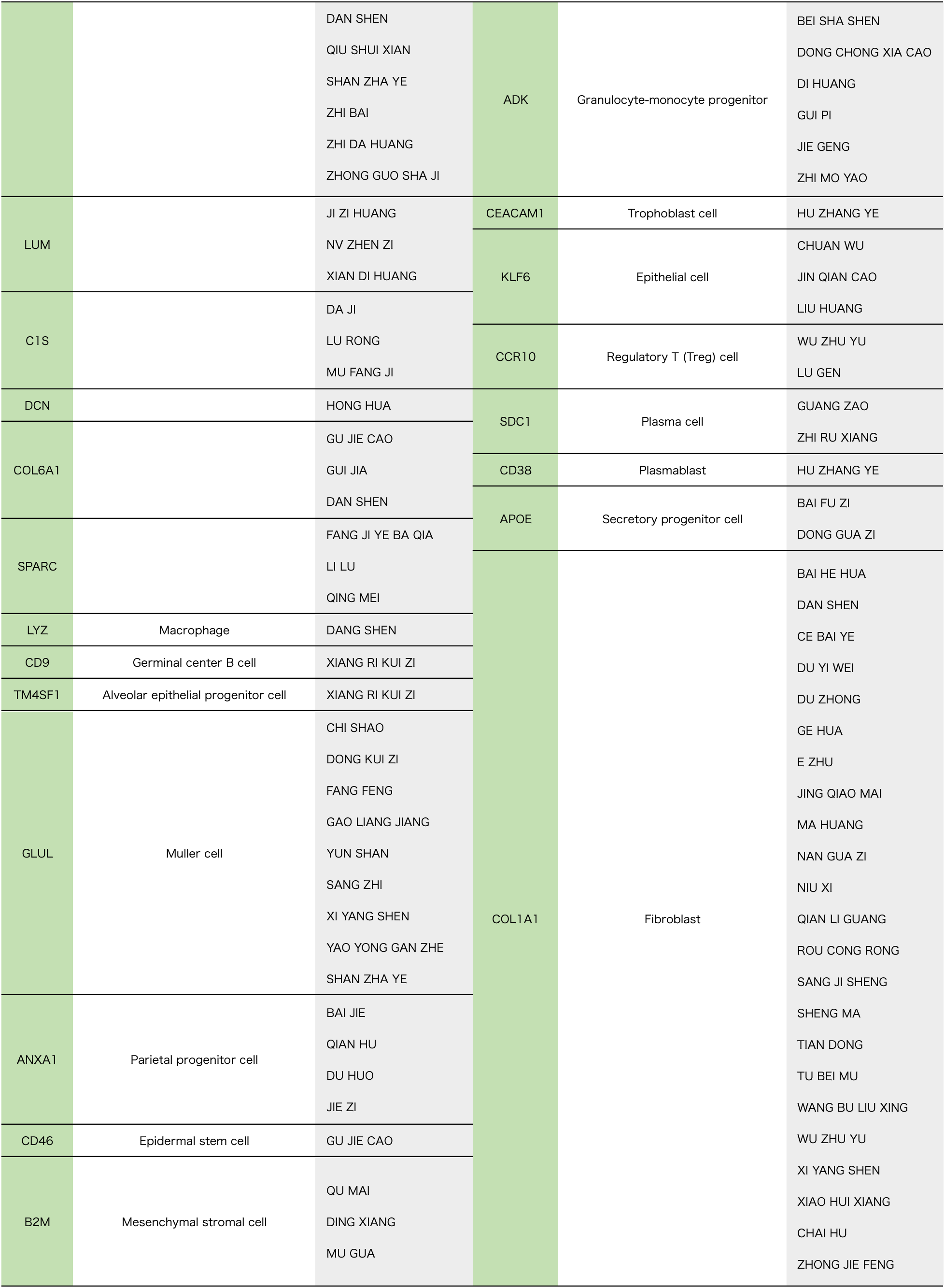
Genes-Cells-Herbs Relationship after Screening with Genes in the TCM Database.

Based on the whole experimental process, it can be speculated that these 8 types of Chinese medicines may regulate the occurrence and development of colorectal cancer by acting on the above-mentioned target genes of these 14 types of immune cells.

### 24 Genes Were Strongly Correlated with 39 Pathways

As shown in Table 3 and Figure 6(a)and(b), we used p-value equal to 0.00E+00 as the filtering condition, and obtained 35 pathways closely related to 39 genes. Most of them were positively correlated, while 12 genes were negatively correlated with the pathways, such as COL1A1 and COL1A2 are negatively to Glycosylphosphatidylinositol GPI anchor biosynthesis; COL1A1, COL1A2, COL3A1, COL6A1 and SPARC were negatively to Biotin metabolism; KLF6 and COL3A1 were negatively to Oxidative phosphorylation; LY86 was negatively to Lysine degradation;COL6A1 was negatively to Terpenoid backbone biosynthesis. COL1A1, COL1A2, COL3A1, COL6A1, LUM, LY86 and SPARC were the respectively high frequency correlations with 35 pathways. In addition, nine pathways had more genes closely related to them, which were Collagen formation, Pantothenate and CoA biosynthesis, Degradation of ECM, Glycosaminoglycan biosynthesis chondroitin sulfate dermatan sulfate, Ferroptosis, IL-10 Anti-inflammatory Signaling Pathway, Inflammatory response, Tumor Inflammation Signature, Apoptosi, Angiogenesis, TGFB and ECM-related gene.

**Figure 6.**
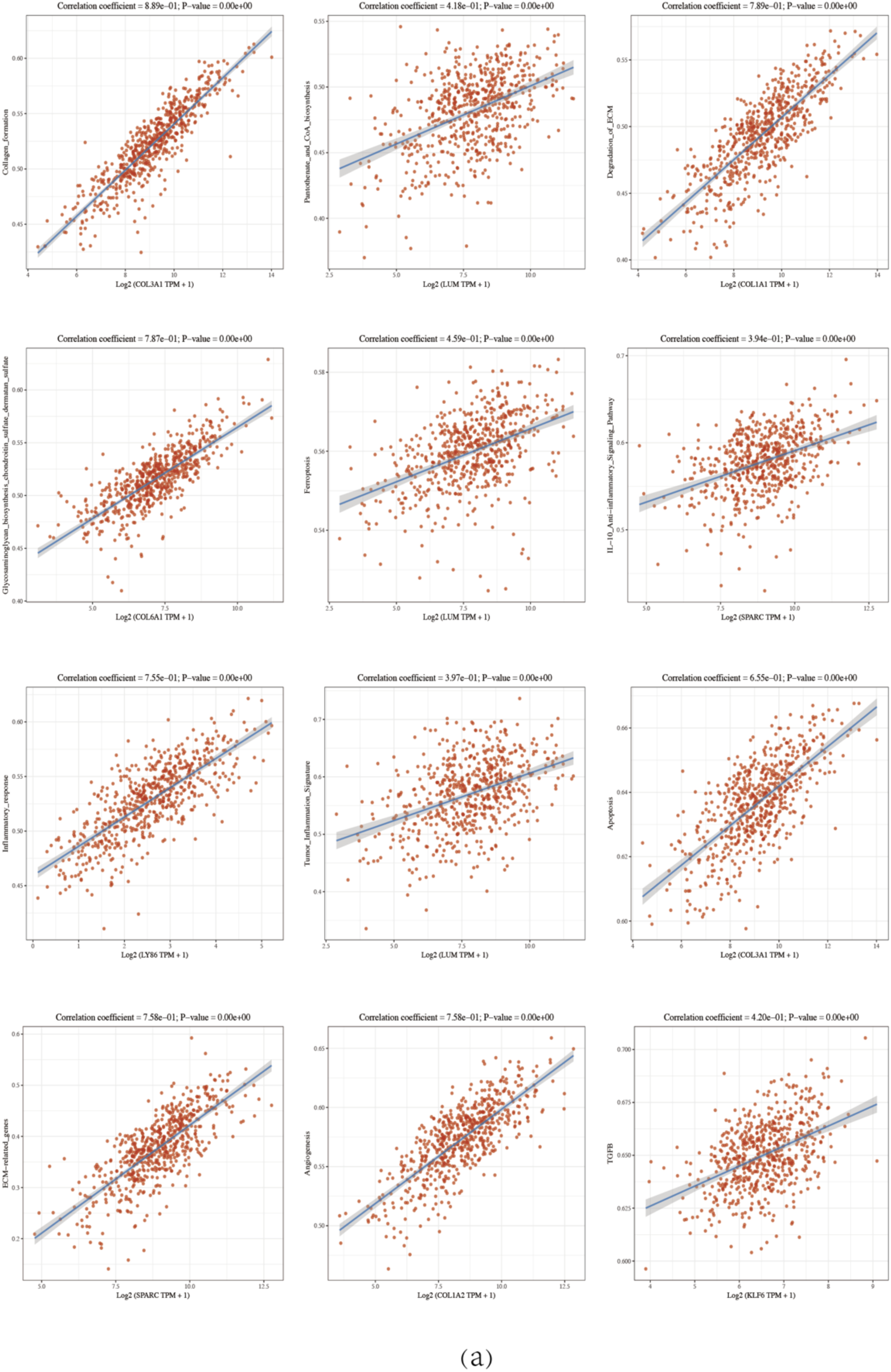
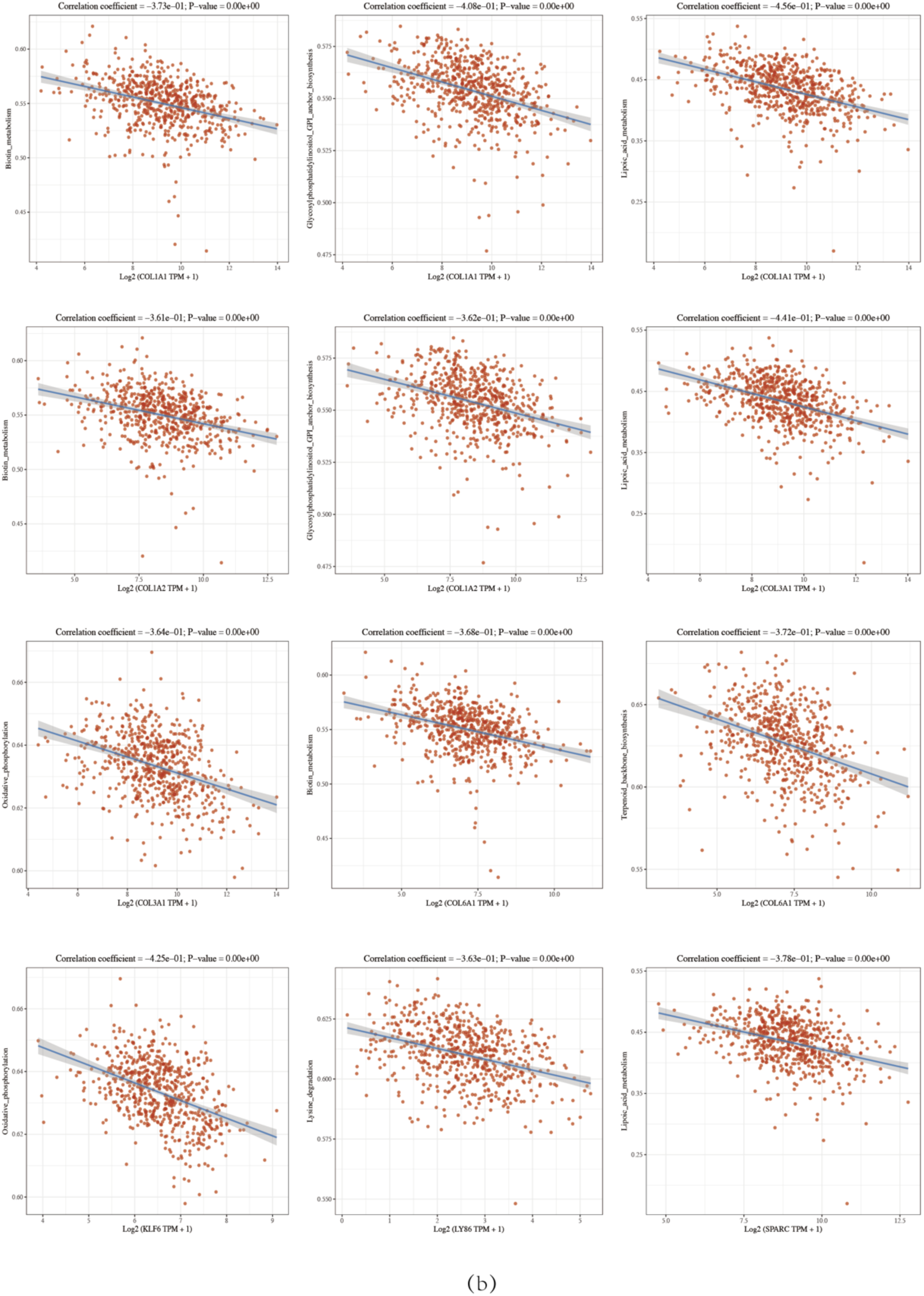
Spearman correlation analysis plots. (a) positive regulation, (b) negative regulation. These Spearman correlation analysis plots are used to show the correlation between the gene expression and the immune score. In these plots, each point represents a sample, and the X-axis and Y-axis represent the gene expression level and immune score, respectively. These are multigene correlation plots, which show the correlation between multiple genes expression and immune scores. In these plots, both the x-axis and y-axis represent genes, and the depth of color represents the size of the correlation coefficient. The deeper the color, the stronger the correlation. The asterisks mark the level of statistical significance, \**p* < 0.05, \*\**p* < 0.01, with the number of asterisks indicating the degree of significance.

**Table 3.**
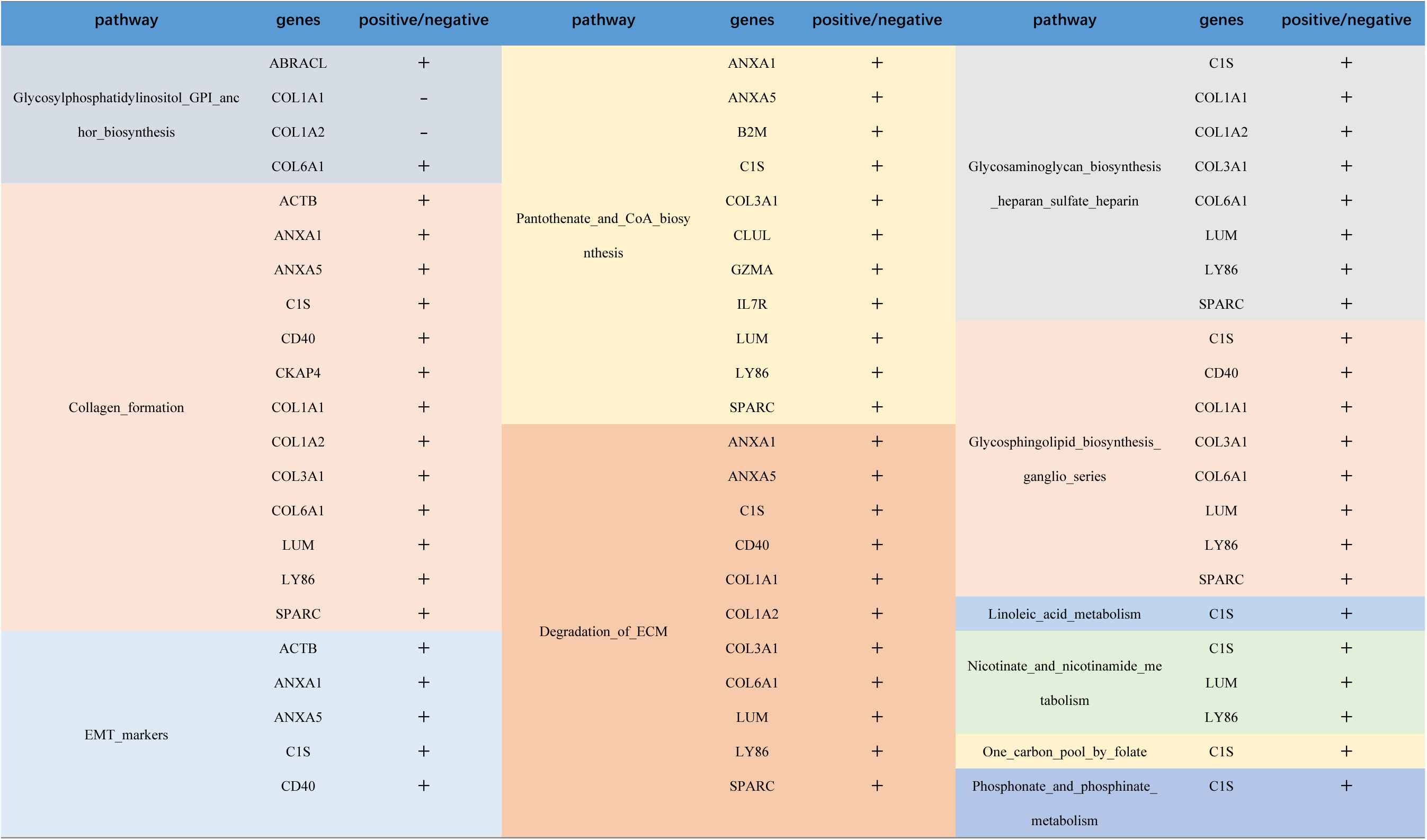
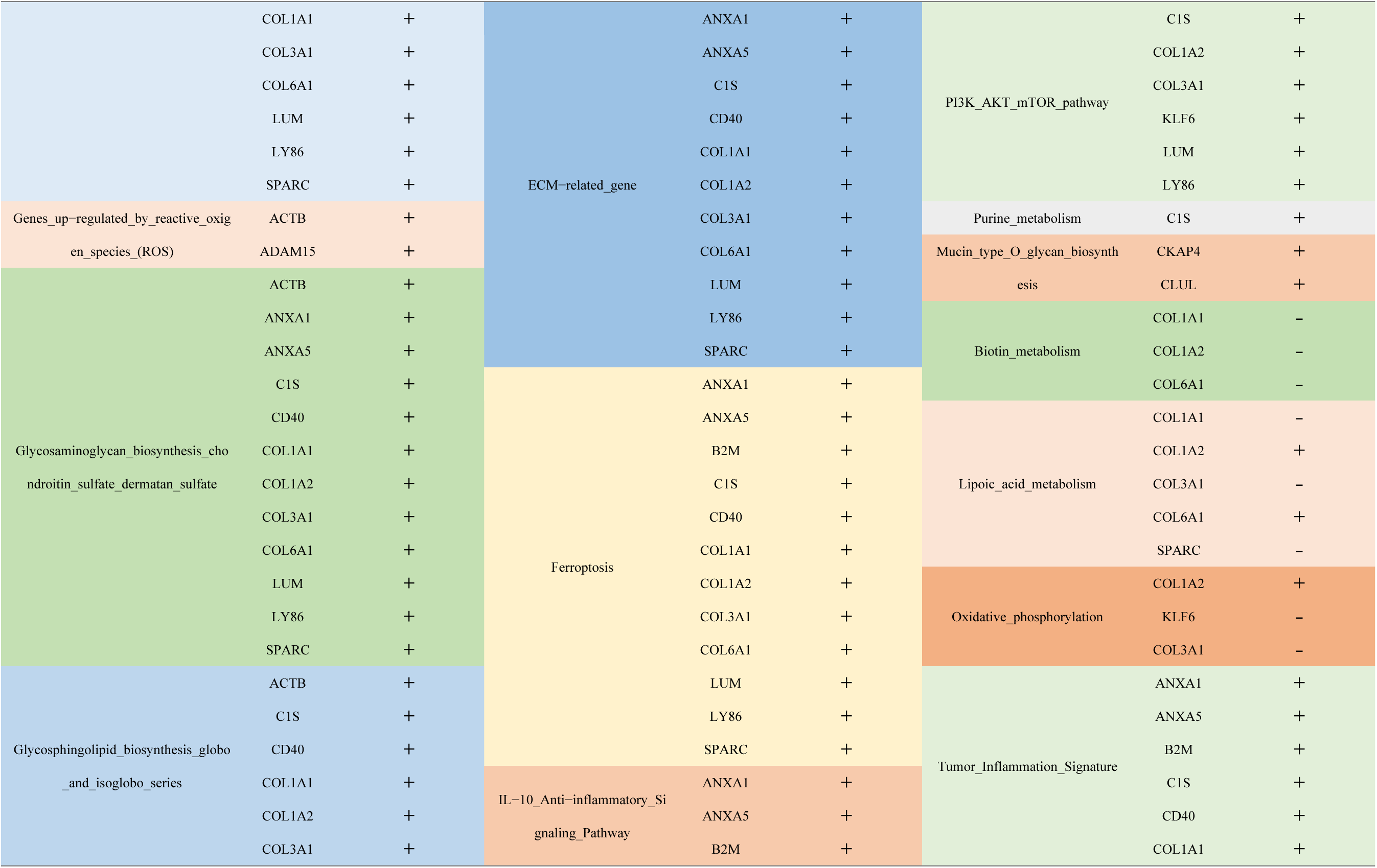
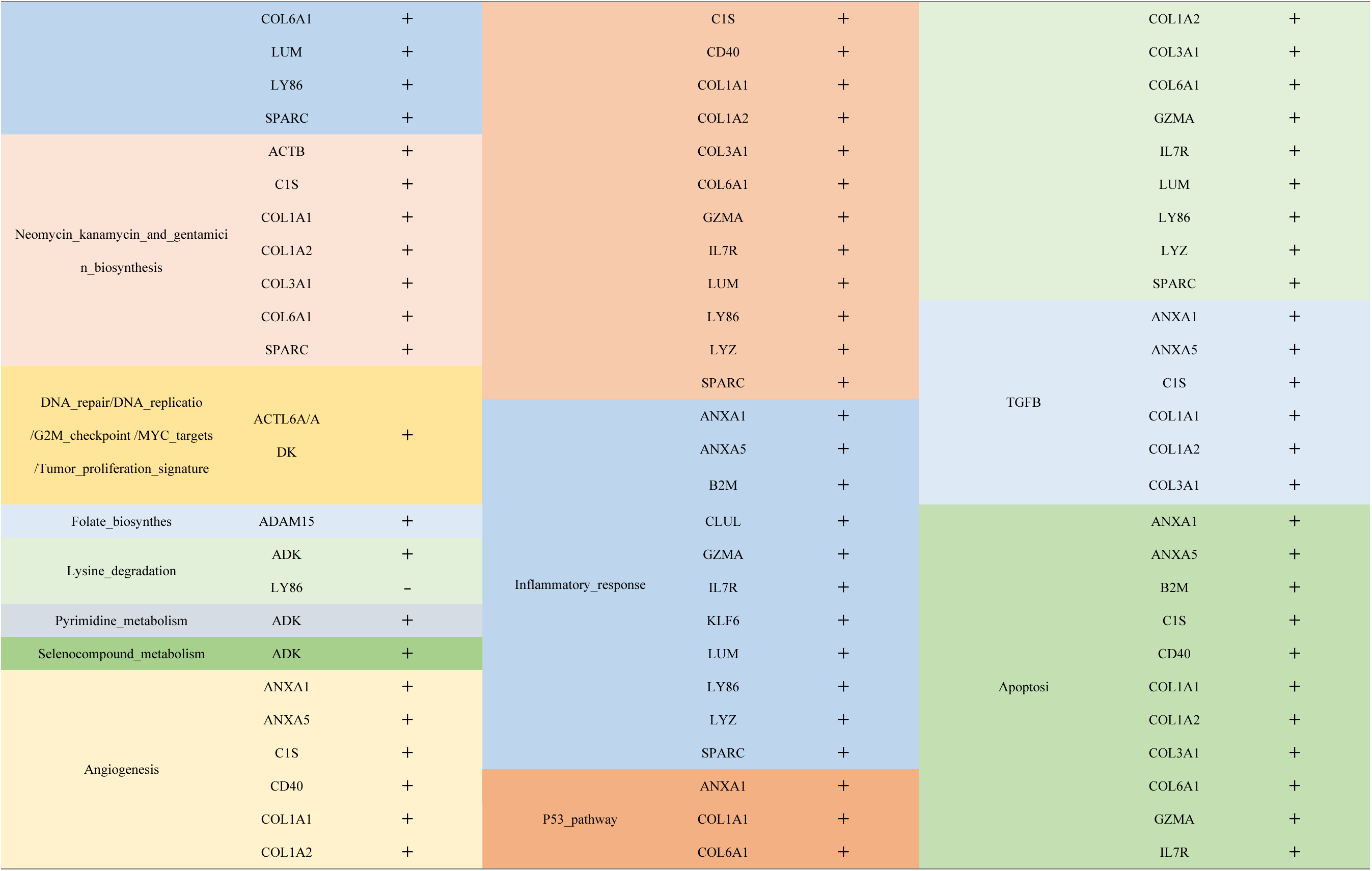
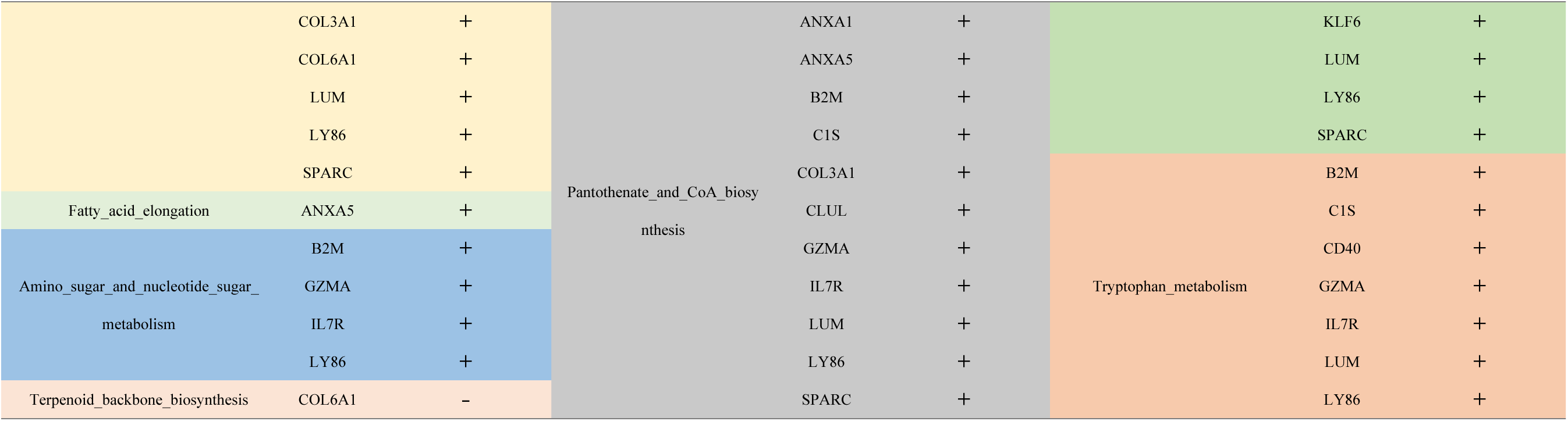
Correlation analysis between genes and pathways using Spearman.

### 23 Genes Associated to Immunity Cells Through 39 Pathways with 6 Algorithms to Analyze

As Figure 7 and Figure 8 shown, for the CIBERSORT algorithm, the 23 genes were strongly associated to main immunity cells and pathways, such as B cells, macrophage M1-M2, mast cells, myeloid dendritic cell activated/resting, NK cell resting, neutrophil, T cell CD4^+^ memory resting, T cell CD8^+^, and T cell regulatory(Treg); In the EPIC algorithm, it was closely associated to main immunity cells, such as B cells, endothelial cells, macrophages, NK cells and T cell CD8^+^,where T cell CD8^+^ might be negative to the most in 23 genes, meanwhile, ADK, ADAM15, ACTL6A, ACTB and ABRACL were also negative to above cells; In the case of MC-counter algorithm, most genes in 23 genes except the lTM4SF1, ADAM15, ACTL6A, ACTB and ABRACL were strongly associated to endothelial cell, macrophage/monocyte, monocyte, myeloid dendritic cell, NK cell, neutrophil, T cell and T cell CD8^+^. Within the quanTiseq algorithm, most genes in 23 were closely associated to B cells, macrophage M1-M2, T cell CD8^+^ and T cell regulatory(Treg). With respect to the TIMER algorithm, it was very significantly for B cells, macrophages, myeloid dendritic cells, neutrophils, T cells CD4^+^, and T cells CD8^+^, where the ADK, ADAM15, ACTL6A, ACTB, and ABRACL genes might negatively regulate the above cells. As far as the xCell algorithm was concerned, it involved a number of immune cells.

**Figure 7.**
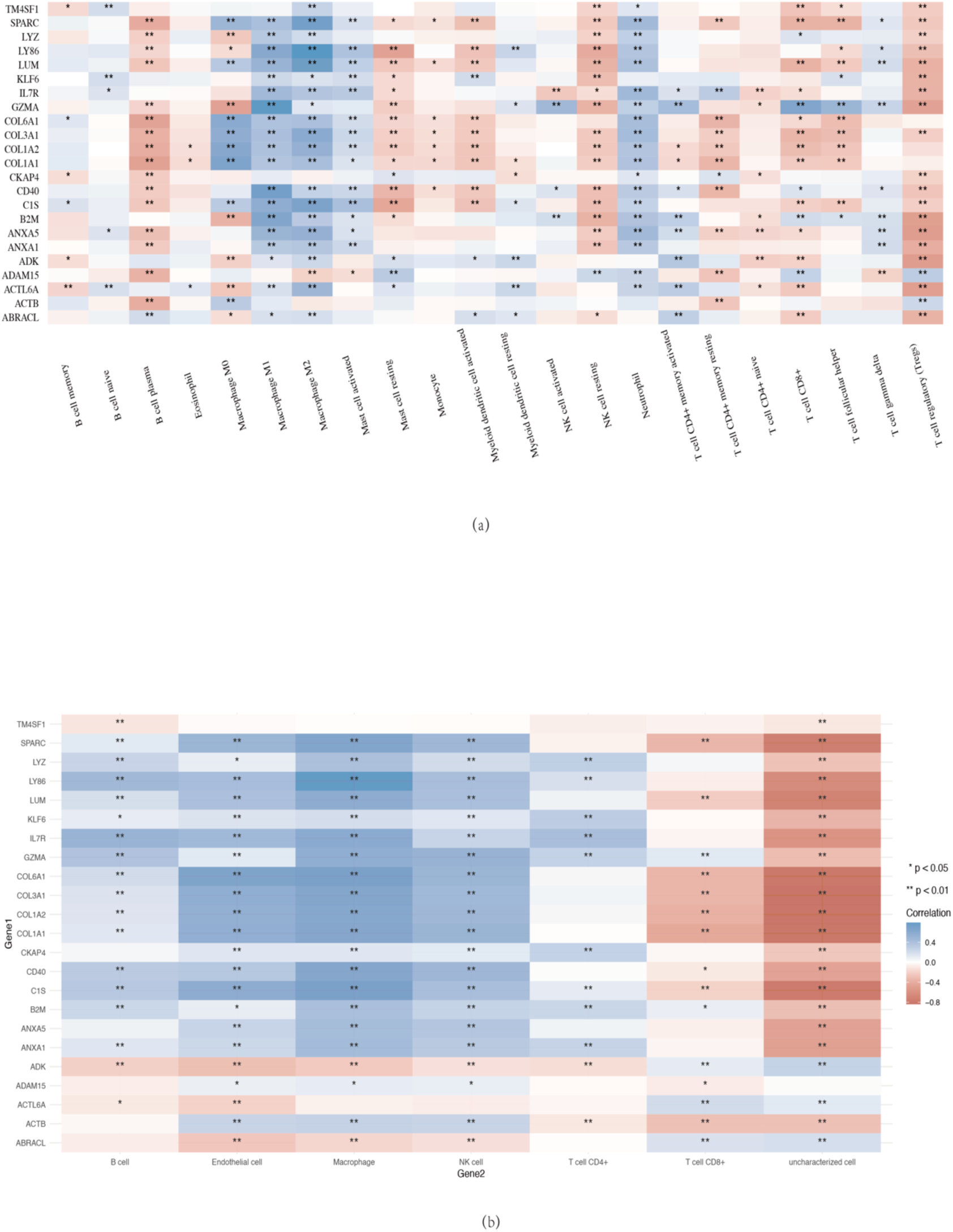
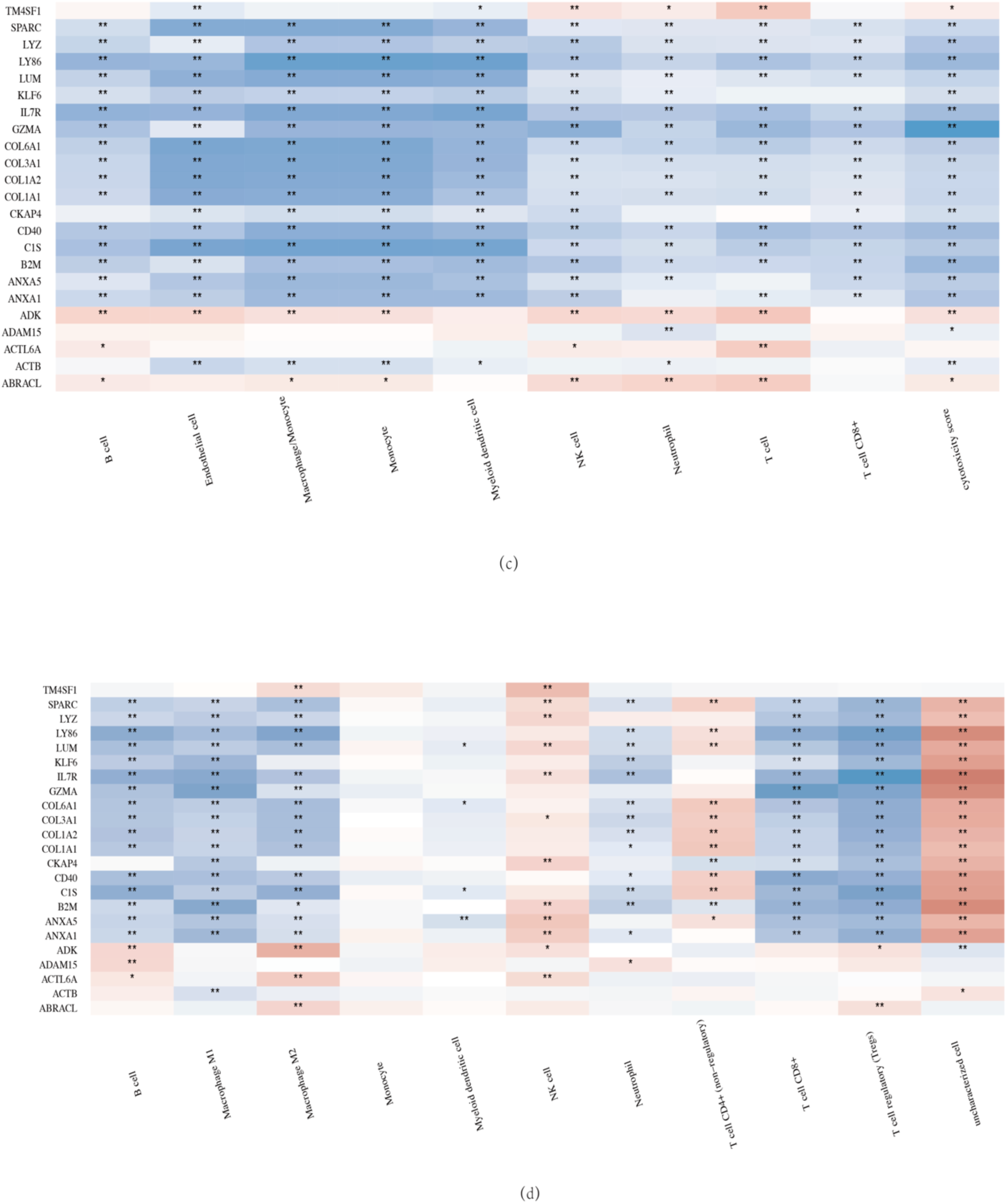
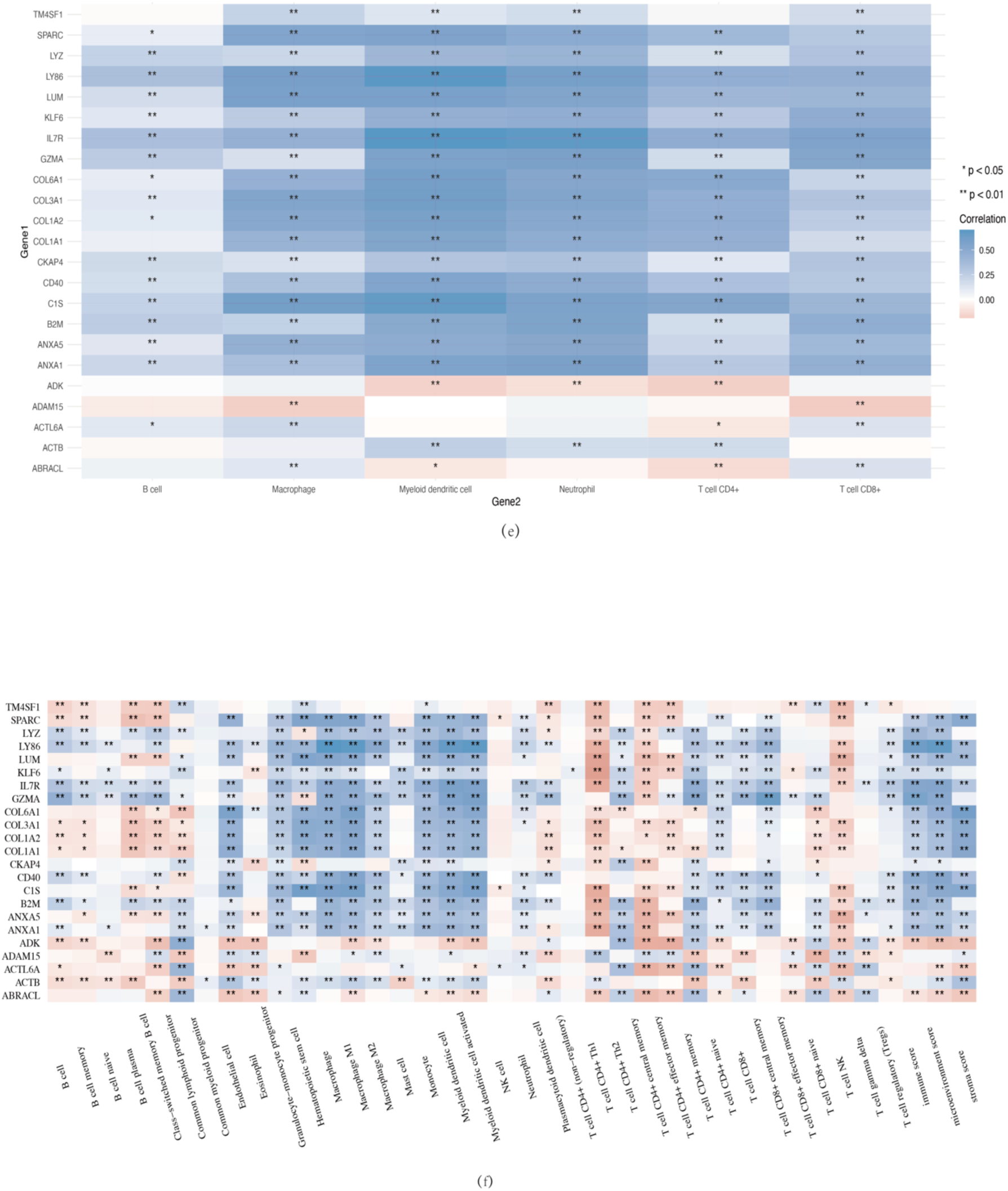
Spearman Correlation Analysis Plots. (a) CIBERSORT algorithm, (b) EPIC algorithm, (c) mcpcounter algorithm, (d) quanTIseq algorithm, (e) TIMER algorithm, (f) xCell algorithm. These Spearman correlation analysis plots are used to show the correlation between the gene expression and the immune score. In these plots, each point represents a sample, and the X-axis and Y-axis represent the gene expression level and immune score, respectively. These are multigene correlation heatmaps, which show the correlation between multiple genes expression and immune scores. In these heatmaps, both the x-axis and y-axis represent genes, and the depth of color represents the size of the correlation coefficient. The deeper the color, the stronger the correlation. The asterisks mark the level of statistical significance, \**p* < 0.05, \*\**p* < 0.01, with the number of asterisks indicating the degree of significance.

**Figure 8.**
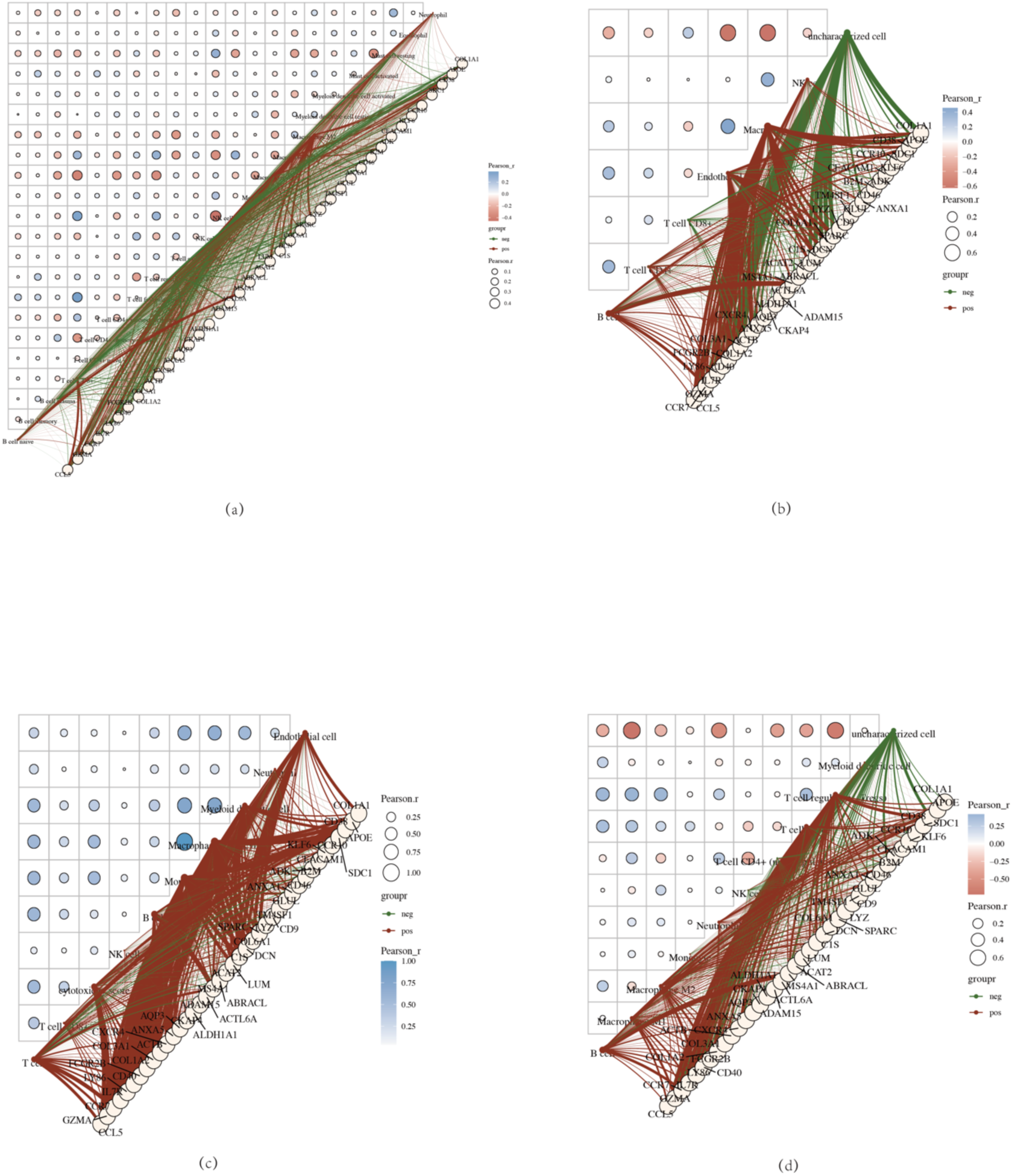
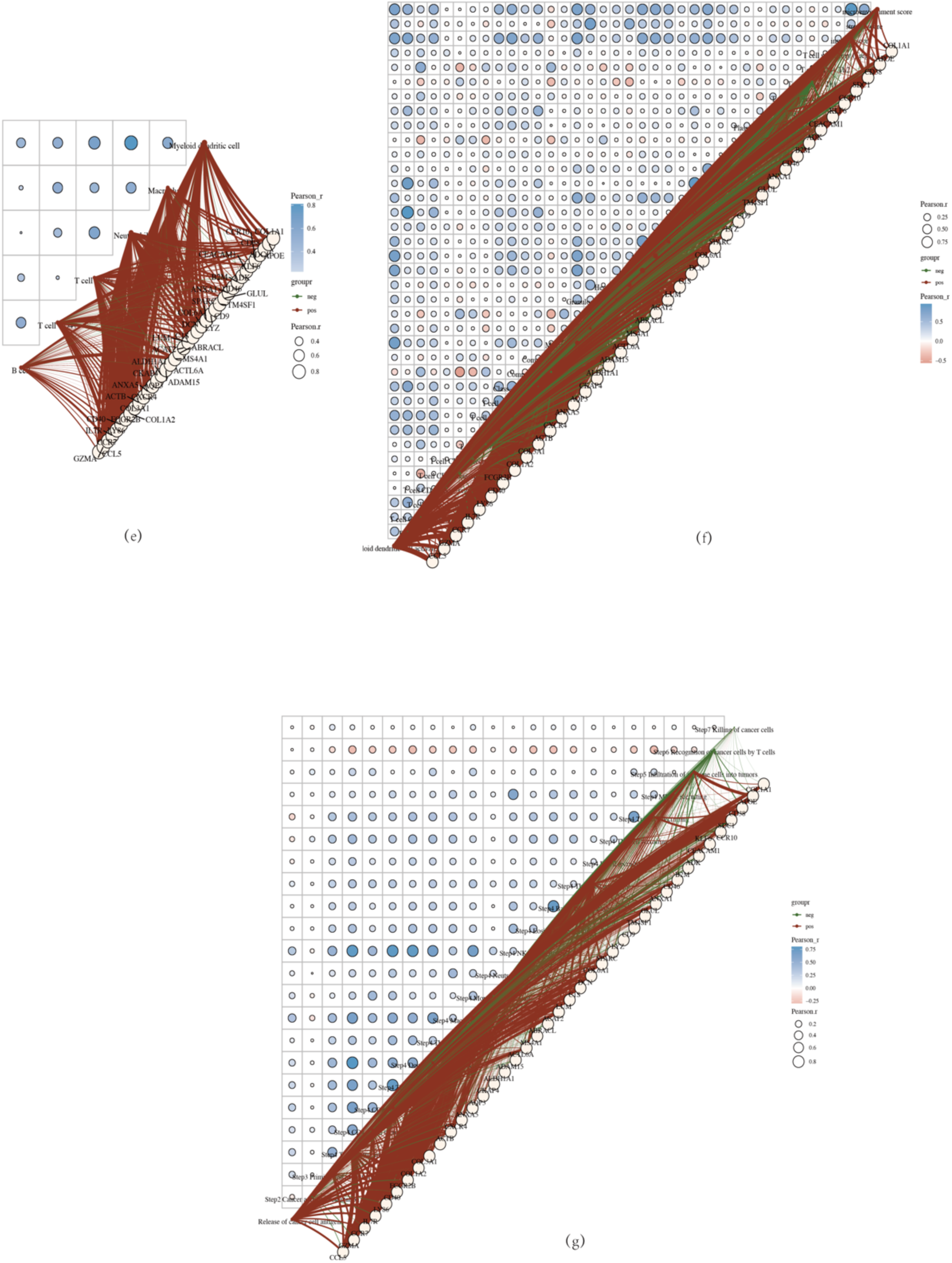
Correlation Analysis Heatmap. (a) CIBERSORT algorithm, (b) EPIC algorithm, (c) mcpcounter algorithm, (d) quanTIseq algorithm, (e) TIMER algorithm, (f) xCell algorithm and (g) TIP algorithm. These represent the correlation analysis among the immune scores themselves, where red represents positive correlation and green represents negative correlation. The redder or greener the color, the greater the correlation between the two; the larger the circle, the stronger the correlation. In the schematic diagram, the red line represents a negative correlation between the model score or gene expression and the immune score, while the green line indicates a positive correlation.

## Conclusion

Single-cell sequencing is a technique for sequencing and analyzing genomic, transcriptomic, and epigenomic levels at the level of a single cell ^(8)^. Traditional sequencing, which is performed on a multicellular basis, actually yields the mean value of the signal in a bunch of cells, losing the information on cellular heterogeneity (differences between cells). The single-cell sequencing technology can detect the heterogeneity information that cannot be obtained by sequencing mixed samples, thus solving this problem well.

Through single-cell analysis, we can not only clarify the differential genes between colorectal cancer tissues and normal tissue cells but also the cell types in which these differential genes are distributed; at the same time, we can annotate the cell functions of these cell types with human cell marker database data^(11)^, so we can conclude that the differential genes of colorectal cancer act on specific cells to perform specific biological functions^(12)^.

Chinese medicine has played a certain role in treating colorectal cancer, especially in reducing the adverse effects of radiotherapy, chemotherapy, targeted therapy, and immunotherapy in colorectal cancer, and its remarkable efficacy^(13)^. However, explaining Chinese medicine’s target and mechanism of action against cancer is difficult because it is a compound preparation with a complex composition. With the establishment of TCM databases in recent years, many TCM databases have focused on the TCM targets, trying to explain the mechanism of action of TCM. Given that single-cell sequencing can focus on specific cells in colorectal tumor cells and can elucidate that certain differential genes in specific cells may be important targets for tumorigenesis and development, combined with the characteristics of the action targets of TCM elucidated in the TCM database, it is easy to closely combine the results of single-cell sequencing and the TCM database to elucidate the action targets of TCM commonly used in TCM clinics for the treatment of colorectal cancer, which is also a novel TCM pathway to elucidate the mechanism of action of TCM.

According to the clustering analysis of the single-cell sequencing results of colorectal cancer tissues, the differential genes in colorectal cancer cells were mainly distributed among fourteen types of cells, and the cell function annotation revealed that most of these fourteen types of cells had immune functions, especially the majority of various types of B cells, stem cells, and T cells. We all know that in the past decade, immunotherapy has achieved impressive success in eradicating malignant cells by harnessing the inherent mechanisms of the host immune system, transforming the therapeutic landscape for a variety of solid and hematological malignancies^(14)^. Immune checkpoint blockade has shown significant benefits. It is increasing the overall survival rates of patients with many different tumors^(15)^. In the treatment of CRC, the PD-1 inhibitors pembrolizumab and nivolumab led a to durable response in patients with metastatic CRC that is mismatch-repair-deficient (dMMR) and microsatellite instability-high (MSI-H) (dMMR-MSI-H)^(16)^. Another inhibitor, ipilimumab, a fully humanized monoclonal antibody that blocks cytotoxic T lymphocyte-associated protein 4, has also been approved by the FDA for combination with nivolumab in patients with dMMR-MSI-H CRC who have previously received chemotherapy^(17)^. Other immunotherapies are currently used against CRC, such as cytokines, cancer vaccines, small molecules, oncolytic viruses, and chimeric antigen receptor therapy ^(18)^. All those immunotherapies have changed the treatment prospects of a variety of solid tumors compared with conventional chemotherapy and targeted therapy, and colorectal cancer is no exception. In a clinic, the use of traditional Chinese medicine in treating colorectal cancer plays a certain therapeutic effect^(3)^. After the analysis of the results of this experiment, anti-tumor Chinese medicine in the human body may be the role of these cells with immune function and play an anti-tumor effect, which suggests that we may be able to get through the single-cell sequencing analysis of the differences in the gene target to, the development of the immunomodulatory role of traditional Chinese medicine anti-tumor drugs.

Chinese medicine treatment of colorectal is based on syndrome differentiation and treatment^(3)^. There are four common types of evidence: colorectal damp-heat type, stasis and toxin accumulation type, spleen and kidney deficiency type, and qi and blood deficiency type^(19)^. Therefore, the Chinese medicines used accordingly have the functions of clearing away heat and removing dampness, resolving blood stasis and detoxifying toxins, strengthening the spleen and the kidneys, tonifying Qi and nourishing blood, and at the same time, they all have the function of eliminating obstruction and accumulation of masses^(20)^. We used the differential genes obtained from single-cell sequencing analysis to screen in the Chinese medicine database and obtained eight classes of Chinese medicines, the functional attributes of which are fully compatible with the needs of clinical diagnosis and treatment in Chinese medicine. For target genes, most of them are similar to the results in the published article through meta-analysis and explore the effective components and potential targets based on the network pharmacology method. It is important to clarify here that we utilized single-cell sequencing specimens sourced from human tumor tissues and that the use of single-cell sequencing technology better excludes clinical mismatches with patients and inaccuracies in differential genes, which more accurately elucidates colorectal cancer genetic variability. For Chinese medicine clinics in treating colorectal cancer, a paper showed in 2017 that a multicenter prospective cohort study enrolled 312 patients with NCCN guidelines including 166 patients who were high exposure to TCM, and follow-up visits were conducted over five years, the result showed that the high exposure to TCM was associated with both better disease-free survival(DFS)(hazard ratio [HR] = 0.62, 95% confidence interval [CI] = 0.39 to 0.98) and overall survival(OS)(HR = 0.31, 95% CI = 0.14 to 0.68)^(21)^. In subgroup exploratory analysis, the effects demonstrated that the differences in outcomes were statistically significant in patients who had received chemotherapy^(21)^. This study concluded that a longer duration of TCM herbal use is associated with improved survival outcomes in part colorectal cancer patients in China. In the face of the problem that the mechanism of action of effective Chinese medicine for the treatment of colorectal cancer is not clear, the results of our study can just be used to reveal its mechanism of action. The results obtained using advanced analytical tools and methods are consistent with the traditional Chinese medicine clinical knowledge of colorectal cancer, which is important to show that it is feasible to analyze the mechanism of Chinese medicine treatment of colorectal cancer by using single-cell sequencing analytical method combined with modern Chinese medicine database.

In recent years, many scholars have put forward the theory of tumor microenvironment (TME) that the growth and metastasis of tumor cells are influenced by their surrounding microenvironment. This microenvironment includes a variety of cells such as fibroblasts, immune and inflammatory cells, and glial cells surrounding the tumor cells, as well as the intercellular matrix, microvasculature, and biomolecules. Understanding the TME has led to the development of novel therapeutic strategies targeting components such as immune checkpoints, angiogenic factors, and stromal cells to inhibit tumor growth and metastasis^(22, 23)^. The results of this study are aptly reflected in the fact that Chinese herbs may play a role in improving the tumor microenvironment by affecting immune cells.

In the results of our analysis, we found that some genes were present at high frequencies, and all of these genes are involved in immune-related processes in groups that have been reported in the literature, such as COL1A1, COL1A2, COL6A1, COL3A1, SPARC, KLF6 and LY86. The protein encoded by the COL1A1 gene is the α1 chain of type I collagen, and there is also a correlation between COL1A1 expression and immune cell infiltration, as evidenced by a positive correlation with the expression of CD4 cells, DC cells, NK cells, macrophages, monocytes, and neutrophils, and a negative correlation with the expression of B cells^(24)^. COL1A2, another subunit of type I collagen, has been found to act as a serum peptide marker for immune-associated pneumonia (CIP), and its expression level is closely associated with the onset and progression of CIP^(25)^. The COL6A1 gene encodes a protein that is the α1 subunit of type VI collagen, may be involved in the regulation of the tumor immune microenvironment^(26)^. The protein encoded by the COL3A1 gene is type III collagen, and platelets interact with type III collagen through specific glycoproteins and non-integrin receptors, suggesting that COL3A1 may be involved in the process of platelet activation and aggregation, which in turn interacts with the immune system^(27)^. SPARC (SPARC-like protein 1) is a protein involved in regulating the synthesis, remodeling and assembly of the extracellular matrix (ECM). In addition to the regulation of the extracellular matrix, SPARC may be involved in biological processes such as inflammation and immune response^(28)^. KLF6 is a tumor suppressor gene that is frequently downregulated in colorectal cancer, and its loss of function is associated with tumor progression, poor differentiation, lymph node metastasis, and advanced TNM stages. KLF6 plays a role in regulating apoptosis, cell cycle, and angiogenesis, making it an important factor in colorectal cancer development and progression^(29)^. LY86 (also known as MD-1) is a protein associated with RP105, which modulates Toll-like receptor 4 (TLR4) signaling in immune cells such as B cells and dendritic cells. It plays an important role in immune regulation by influencing B cell activation and dendritic cell function, but it does not directly bind to or interact with TLRs. Instead, the RP105/MD-1 complex indirectly affects TLR4 signaling^(30)^. All of these genes were the emergence of our strict screening conditions (*p*-value equal to 0.00E+00), therefore, it can be presumed that for colorectal cancer plays a major role in the immune process, but also it is not difficult to speculate that traditional Chinese medicine may also act on the process of the expression of the above genes and control the process of tumor development, progression and healing.

In our study, we also obtained several pathways closely related to the above genes, a dozen of which are positively and negatively correlated. Through the collagen-forming pathway, collagen molecules also contain immunoreactive molecules, which are involved in the immune regulation of the body. In addition, collagen degradation products may act as “red flags” for the immune system, triggering an immune response^(31)^. For pantothenate and CoA biosynthesis, whose abnormal biosynthesis may be associated with certain immune-related diseases. In addition, CoA is involved in the β-oxidation of fatty acids, which provides energy to immune cells. In the case of degradation of ECM, the fragments and signaling molecules generated during degradation can activate the immune system and trigger an immune response. Also, immune cells participate in the degradation process of the extracellular matrix by secreting enzymes^(32)^. In terms of glycosaminoglycan biosynthesis, which can be involved in the recruitment, activation, and migration processes of immune cells, affecting the intensity and duration of the immune response^(33)^. For the well-known ferroptosis, the oxidative stress and inflammatory response generated during iron death can activate the immune system and trigger an immune response. At the same time, immune cells also influence the process of iron death by regulating iron ion metabolism and lipid peroxidation^(34)^. Regarding the IL-10 Anti-inflammatory Signaling Pathway, IL-10 is an important anti-inflammatory cytokine. activation of IL-10 anti-inflammatory signaling pathway can inhibit the activation and proliferation of immune cells, and reduce the inflammatory response and tissue damage^(35)^. With the respect to Inflammatory response, immune cells participate in the inflammatory response process by recognizing, phagocytosing, and removing pathogens or damaged cells. At the same time, signaling molecules such as cytokines and chemokines produced during the inflammatory response can also regulate the activation and migration process of immune cells^(36)^. As for tumor inflammation signature, tumor inflammation signature is an important part of the tumor microenvironment, and is closely related to the processes of tumor proliferation, invasion and metastasis. Immune cells trigger immune responses by recognizing antigens on the surface of tumor cells or tumor-associated antigens, which in turn are involved in the immunotherapy of tumors and the development of immunotherapy resistance^(37)^. For apoptosis, immune cells remove apoptotic cells by recognizing signaling molecules on the surface of apoptotic cells to prevent inflammatory reactions and autoimmune diseases. At the same time, apoptotic cells can also recruit immune cells for clearance by releasing “eat me” signals^(38)^. When it comes to angiogenesis, or blood vessel angiogenesis, is a key component of tumor growth and metastasis. Immune cells regulate tumor angiogenesis by secreting angiogenic factors or factors that inhibit angiogenesis. Meanwhile, microenvironmental changes such as hypoxia and nutritional deficiencies generated during tumor angiogenesis can also affect the activation and function of immune cells^(39)^. TGFB (transforming growth factor β) and ECM-related genes are involved in a variety of biological processes, including cell proliferation, differentiation, apoptosis and immune regulation^(40)^. Extracellular matrix-related genes, on the other hand, encode molecules such as proteins and polysaccharides that make up the extracellular matrix. These molecules influence the onset and development of immune responses by regulating the adhesion and migration processes of immune cells. Meanwhile, abnormal expression of TGFB and extracellular matrix-related genes may also be associated with the development of certain immune-related diseases. Refer to biotin metabolism, biotin deficiency leads to decreased immune system function and increased risk of infection^(40)^. Glycosylphosphatidylinositol (GPI) anchor biosynthesis, in the immune system, GPI-anchored proteins play key roles in B-cells, T-cells, and a variety of immune receptors. They are involved in the regulation of immune response by recognizing and binding to the corresponding ligands, triggering the activation, proliferation and differentiation of immune cells^(41)^. As for oxidative phosphorylation, it plays an important role in the regulation of macrophage function and polarization, which in turn affects the intensity and type of immune response^(42)^. Terpenoid backbone biosynthesis have received much attention in biosynthetic herbal medicine-related studies;Terpenoids are a class of natural products with a wide range of biological activities in the immune system and have significant immunomodulatory effects. Some terpenoids inhibit the proliferation and activation of immune cells, thereby reducing inflammatory responses, while others promote the proliferation and differentiation of immune cells and enhance immune responses. In addition, terpenoids can act as signaling molecules involved in the signaling process of immune cells. Therefore, terpenoid skeleton biosynthesis is important for maintaining the balance and homeostasis of the immune system^(43)^.

For the gene-immunity correlation analysis, we analyzed the gene-immunity correlation through six algorithms, and finally found that 23 genes have extremely strong correlation with immune cells, some of which play a positive adjustment role, and some of which play a negative regulation role, such as B cells, macrophage M1-M2, mast cells, myeloid dendritic cell activated/resting, NK cell resting, neutrophil, T cell CD4^+^ memory resting, T cell CD8^+^, and T cell regulatory(Treg); All of these cells have been extensively documented to play a role in tumorigenesis, progression and prognosis^(44, 45)^. We hypothesize that the positive and negative regulatory effects of these genes acting on immune cells may have analogies to the Yin and Yang regulatory theory in Chinese medicine theory.

The TCM doctrine of constitution has a similar viewpoint. According to the theory of TCM, human body is an organic whole, and an imbalance of Yin and Yang in the organism will form a constitution suitable for tumor development. This imbalance may be due primarily to the TCM pathogenesis of cancer as phlegm, dampness, stasis, heat, and toxicity^(46)^. By adjusting the balance of Yin and Yang in the human body, TCM can change the constitution of tumor growth, thus inhibiting the occurrence and development of tumor^(47)^. It may be possible for colorectal cancer by using the 8 types herbs mentioned in this study for clearing heat, clearing away toxins, relieving dampness, removing blood stasis, strengthening the spleen, benefiting the kidney, tonifying Qi, and nourishing the blood to regulate the patient’s righteousness, achieve harmony of Yin and Yang, and maintain internal equilibrium. There is no concept of immunity in TCM, but there is a concept of righteousness and a saying “maintain righteousness within, and evil cannot interfere”^(48)^. The author also believes that the tumor microenvironment here is equivalent to the constitution of a tumor patient in TCM.

In summary, based on our results, we propose that clinical TCM treatment of colorectal cancer may regulate the occurrence and development of colorectal cancer through the action of TCM on immune cells to regulate immunity based on single-cell sequencing analysis combined with TCM database screening. This may be a novel approach to explain the mechanism of action of TCM in the treatment of colorectal cancer.

## Consent for Publication

Not applicable.

## Availability of Data and Materia

We obtained the raw data of single-cell transcriptome profiling(Tissue: colorectal cancer, Data Set Number: GSE163974, Chip Platform: GPL16791, Sample Type: 3 normals and 3 tumors, Species: human, Data Types: 10x Genomics) from the Gene Expression Omnibus (GEO) database (see: https://www.ncbi.nlm.nih.gov/geo/ for more details). The Chinese Herbs described in the manuscript were screened on the database named HERB (A high-throughput experiment-and reference-guided database of traditional Chinese medicine) (see: herb.ac.cn/ for more details). The STAR-counts data obtained in the immune correlation analysis and the corresponding clinical information were downloaded from (https://portal.gdc.cancer.gov).

## Specific Author Contributions

Chenlong Zhang works as first author; Concept and design: Chenlong Zhang, Yumei Zhang; Acquisition of data: Chenlong Zhang, Yumei Zhang, Yujie Wang, Kaihang Guo, Pengfei Li, and Chunfang Zhang; Statistical analysis and interpretation of data: Chenlong Zhang and Yumei Zhang; Drafting and revision of the manuscript: Chenlong Zhang, Yumei Zhang, and Chunfang Zhang.

## Acknowledgments

We sincerely thank all the clinical staff at Xiang’an Hospital of Xiamen University for their contributions to this work.

